# STDP forms associations between memory traces in networks of spiking neurons

**DOI:** 10.1101/188938

**Authors:** Christoph Pokorny, Matias J. Ison, Arjun Rao, Robert Legenstein, Christos Papadimitriou, Wolfgang Maass

## Abstract

Memory traces and associations between them are fundamental for cognitive brain function. Neuron recordings suggest that distributed assemblies of neurons in the brain serve as memory traces for spatial information, real-world items, and concepts. However, there is conflicting evidence regarding neural codes for associated memory traces. Some studies suggest the emergence of overlaps between assemblies during an association, while others suggest that the assemblies themselves remain largely unchanged and new assemblies emerge as neural codes for associated memory items. Here we study the emergence of neural codes for associated memory items in a generic computational model of recurrent networks of spiking neurons with a data-constrained rule for spike-timing-dependent plasticity (STDP). The model depends critically on two parameters, which control the excitability of neurons and the scale of initial synaptic weights. By modifying these two parameters, the model can reproduce both experimental data from the human brain on the fast formation of associations through emergent overlaps between assemblies, and rodent data where new neurons are recruited to encode the associated memories. Hence our findings suggest that the brain can use both of these two neural codes for associations, and dynamically switch between them during consolidation.

## 1. Introduction

The formation of associations between memory traces is a fundamental operation in cognitive computations of the brain. However, whereas there is common agreement about the role of assemblies of neurons as neural codes for memory items (Buzsáki 2010, Josselyn et al. 2015, Quian Quiroga 2016), there are conflicting experimental data and models regarding neural codes for associated memory items.

An overview of models for the formation of associations in neural networks can be found in (Kahana et al. 2008, Kahana 2012). Two types of models can be distinguished: chaining models and hierarchical models. A chaining model postulates that synaptic connections between the assemblies for two memory items are enhanced when both items become associated, and that none or only few additional neurons are recruited for representing the combination of the two memory items. In contrast, in a hierarchical model the combination of two memory items is encoded by a new set of neurons that do not belong to the assembly codes for the individual memory items, see e.g. (Norman and O’Reilly 2003).

Experimental support for the hierarchical model was provided by (Komorowski et al. 2009), where neurons in the rodent brain were found that responded to a combinations of an object and a place, but not to the individual object or place. However only associations between items from these two categories, objects and places, were studied in (Komorowski et al. 2009). These associations resulted from extensive training, rather than from associations that were formed “on the fly”.

The rapid formation of associations between two images, or rather the concepts which they evoke, were studied with in vivo single-neuron recordings in human epilepsy patients implanted, for clinical reasons, with depth electrodes in the Medial Temporal Lobe (MTL) (Ison et al. 2015, De Falco et al. 2016). These studies found that very few neurons in the MTL responded only to a combined image, but not to any of its two components. Instead, their data suggest that an association between two memory items is encoded through a modification of the assemblies of neurons that encode the two memory items involved: Each of them expands and recruits neurons from the other assembly. Hence these data support a chaining model for the formation of associations.

We investigated the question which of these conflicting models and experimental data for the formation of associations could be reproduced by a generic model for a recurrent network of neurons in the brain, with a data-constrained rule for synaptic plasticity. In order to make the dynamics of the model similar to that of networks of neurons in the brain, we used standard models for excitatory and inhibitory spiking neurons, and included data-constrained short-term synaptic plasticity, i.e., individual mixtures of synaptic facilitation and depression that vary according to (Gupta et al. 2000) with the type (excitatory or inhibitory) of the pre- and postsynaptic neuron. We modelled longterm synaptic plasticity with a data-constrained rule for Spike-Timing Dependent Plasticity (STDP), the triplet rule (Pfister and Gerstner 2006). This rule can reproduce a variety of experiments, such as pairing experiments with frequency effects (Sjöström et al. 2001), and triplet and quadruplet experiments (Wang et al. 2005). We show that the resulting neural network model can reproduce the data on emergent neural codes for associations in the human MTL from (Ison et al. 2015, De Falco et al. 2016). In fact, several details of the experimental data of (Ison et al. 2015) are reproduced by the model. On the other hand, we find that a different parameter setting for the same neural network reproduces the neural code that is postulated by hierarchical models, where a new assembly of neurons is assumed to become the memory trace for the combined memory. Hence our results suggest that the brain is able to generate both types of neural codes for combined memories. It is even possible that the same neural network in the brain switches between these two different representations of associated memories in the course of consolidation.

By making explicit the conditions under which memory traces and different types of neural codes for associations can be reproduced by a generic neural network model, these results pave the way for larger memory models that enable an analysis of the complex web of associations in the brain.

## 2. Methods

### 2.1 Details of the network model

Our network model is a generic recurrent network of spiking neurons. The details of the results of our simulations depend on the specific parameter settings, such as the specific realization of the synaptic connectivity matrix that is drawn according to given connection probabilities, the choice of initial values of synaptic weights, delays, and short-term synaptic plasticity from given distributions, and on the specific realization of rate patterns that serve as inputs to the network. But the main findings were independent of these specific choices.

In the following subsections, the default setting of parameter values that were used in our model can be found. They are referred to as the *standard values*. We refer to one specific randomly drawn realization of the network based on these standard values, together with a fixed triple of input patterns (see Section 2.2.1), as the *standard model*. Specifically, the results that are reported in Figures 3 to 6, and S5 in the *Supplementary Material* are based on this standard model. We additionally analyzed in Figures 5C-E and 6 how the results depended on the specific realization of the network and input patterns by randomly generating different networks (with newly drawn connectivity and initial parameters) and input pattern triples.

**Figure 1:**
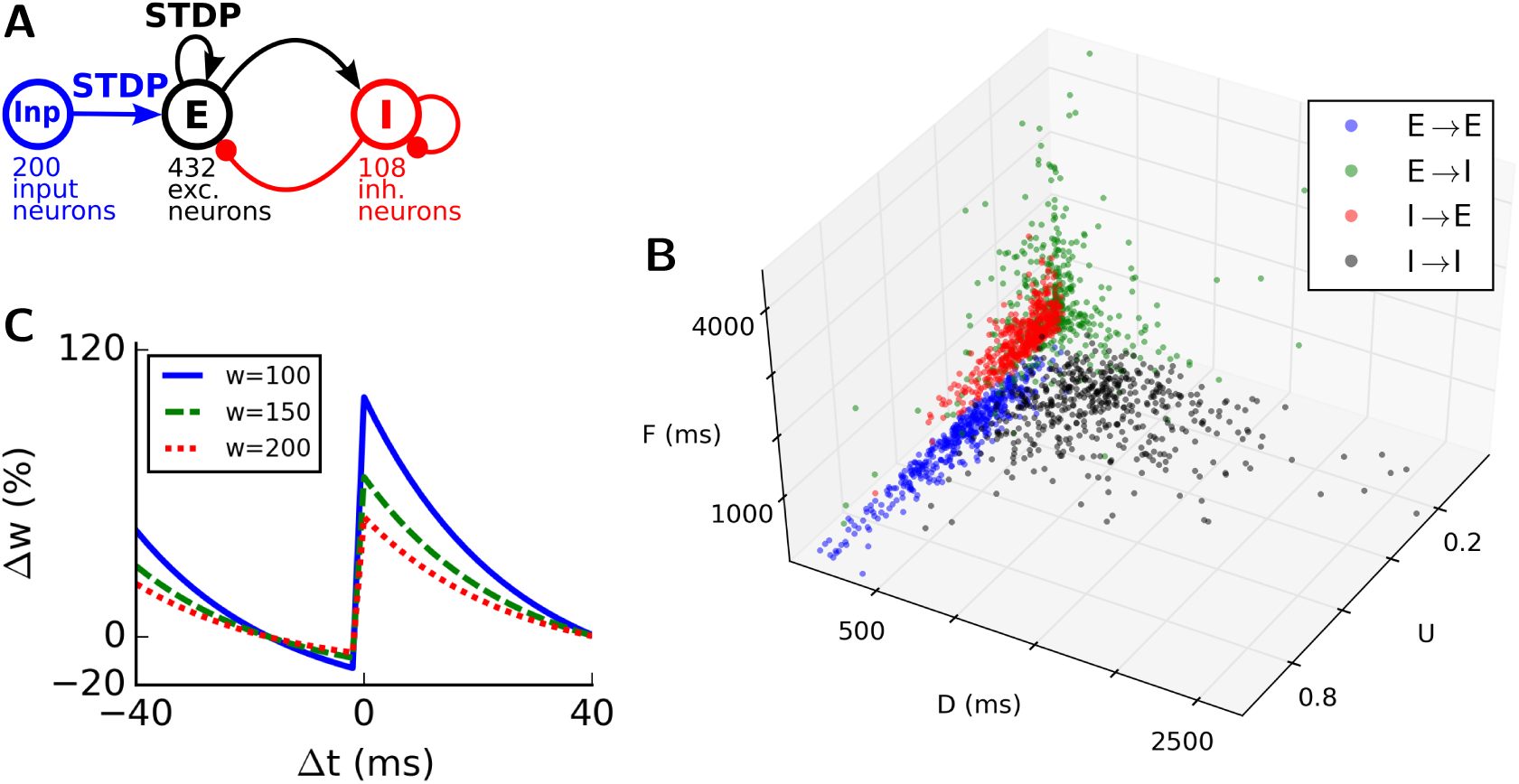
(A) Network architecture, consisting of 200 input neurons (Inp), 432 excitatory neurons (E), and 108 inhibitory neurons (I). Inp*→*E as well as recurrent E*→*E synapses were subject to STDP. (B) Illustration of the distribution of UDF parameter values for short-term synapse dynamics in three-dimensional parameter space for different types of connections (blue: E*→*E; green: E*→*I; red: I*→*E; black: I*→*I). UDF values were generated from bounded gamma distributions with means and SDs given in Table 2. (C) STDP triplet rule that was used in our model for synapses from external inputs and recurrent excitatory synapses, showing the relative weight change after 10 spike pairings at a pairing frequency of 20 Hz for different initial weight values.

**Figure 2:**
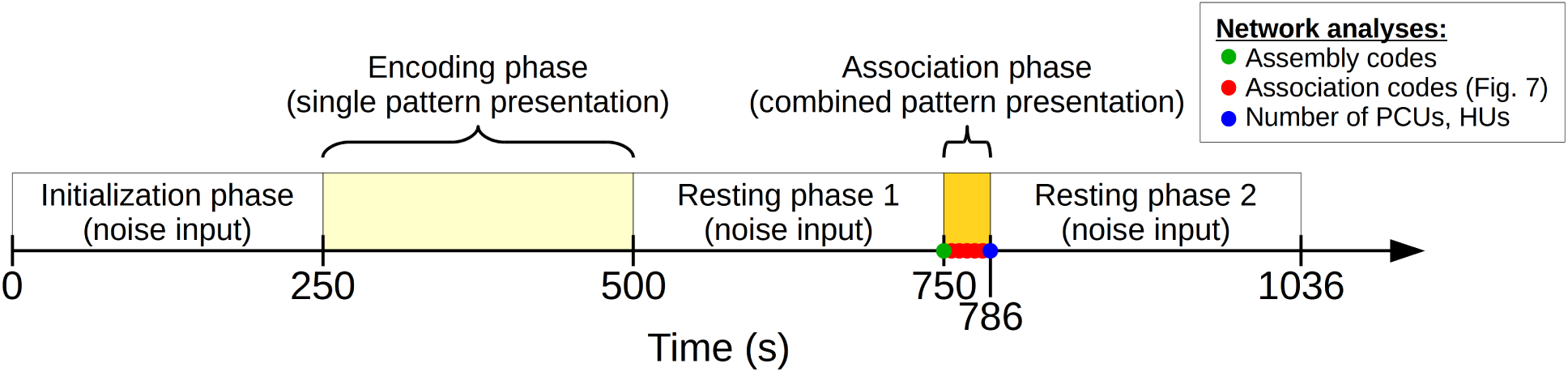
Scheme of the experimental setup, showing the five input phases of the main simulation during which different kinds of input patterns or noise was presented as input to the network. The network was analyzed at time points indicated by circular markers (green: analysis to determine assembly codes, see Section 2.2.4; red: stepwise analyses to determine association codes for neuronal and functional learning curves, see Section 2.4; blue: analysis to determine numbers of pair-coding units (PCUs) and hierarchical units (HUs), see Sections 2.3.2 and 2.3.4).

**Figure 3:**
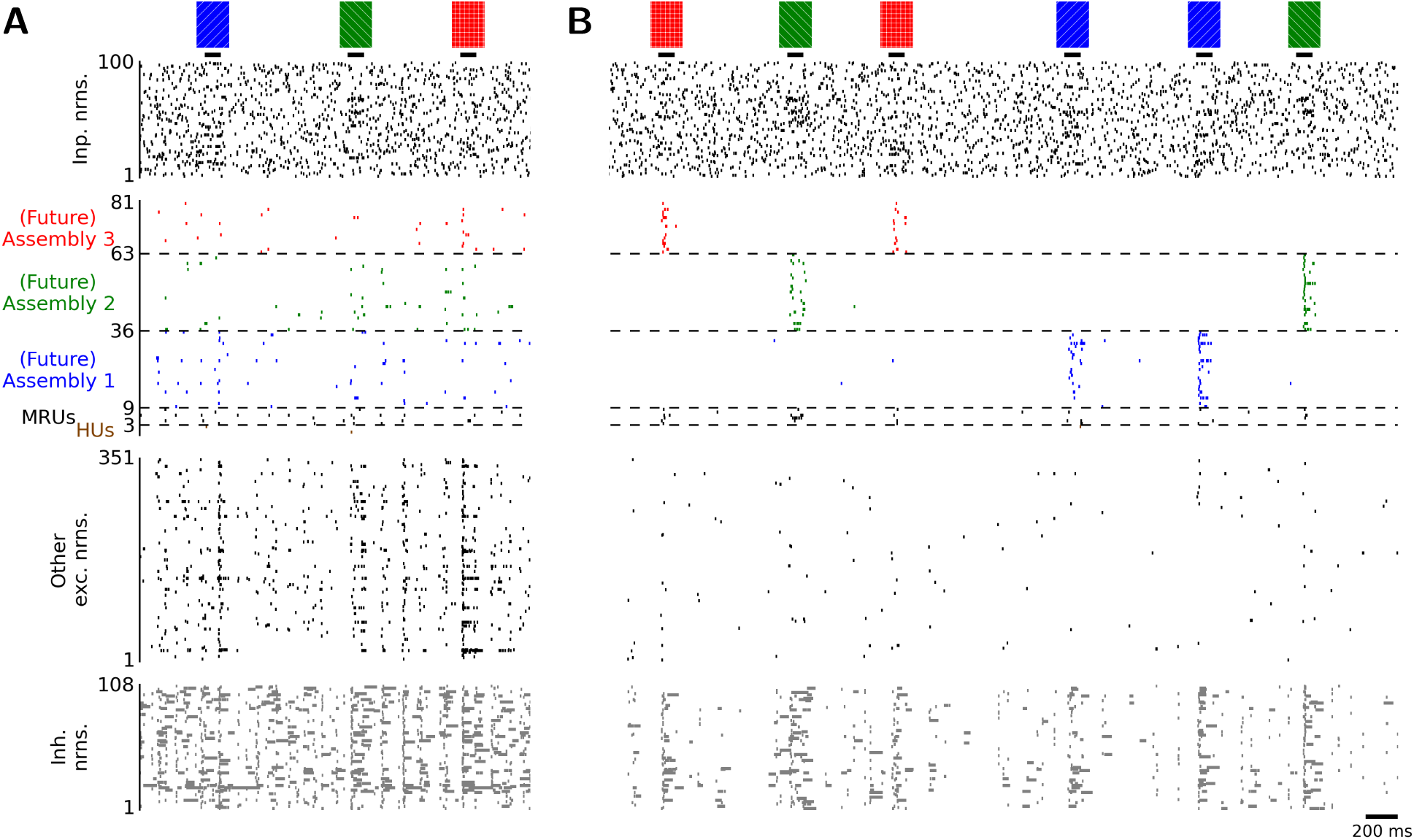
Emergence of assembly codes for repeated input patterns. (A) Initial network activity in response to the three input patterns, before any synaptic plasticity took place. The excitatory neurons are grouped into the assemblies to which they will belong after the encoding and subsequent resting phase 1. (B) Emergence of assemblies after the encoding and subsequent resting phase 1. Neurons in the three assemblies responded preferentially to one of the three input patterns. Spike trains are shown (from top to bottom) for input neurons (only the first 100 of 200 are shown), assembly neurons, multi-responsive units (MRUs, responding to more than one input pattern), (future) hierarchical units (HUs), other excitatory neurons, and inhibitory neurons. Note the different y-axis scalings to highlight neurons in assemblies, MRUs, and HUs.

**Figure 4:**
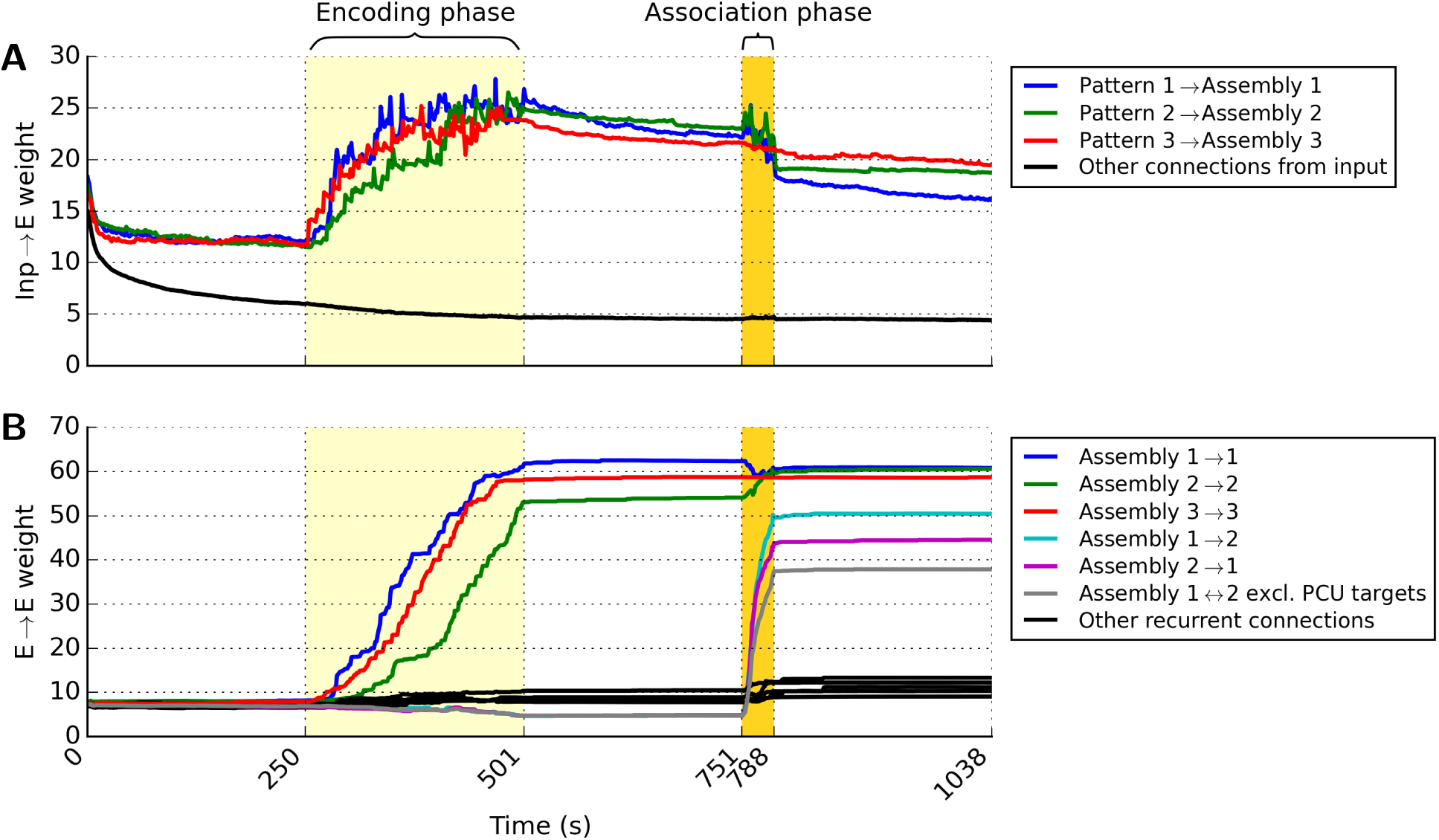
Mean weight changes of (A) connections from input neurons to the three assemblies, taking only on-channels of the stationary input rate patterns into account (see text) and (B) internal connections within the three assemblies during all phases of the main simulation. The following 5 phases were analyzed, which were defined by the external input: (1) Initialization phase of 250 s with just noise input, (2) encoding phase of 250 s with three input patterns randomly presented to the network, (3) resting phase 1 of 250 s with just noise input, (4) association phase of 36 s with a combined pattern repeatedly presented to the network, and (5) resting phase 2 of 250 s with just noise input. The input neurons fired with a baseline rate of 5 Hz during the 3 resting phases. During all 5 phases, both Inp*→*E and E*→*E synapses were subject to STDP. Colored lines show mean weights over time as defined in the legends. Black lines denote mean weight changes of all other synaptic connections between excitatory neurons, i.e., cross-connections between other pairs of assemblies and connections not related to any assembly. The gray line shows that also weights to non-PCUs increase during the association phase.

**Figure 5:**
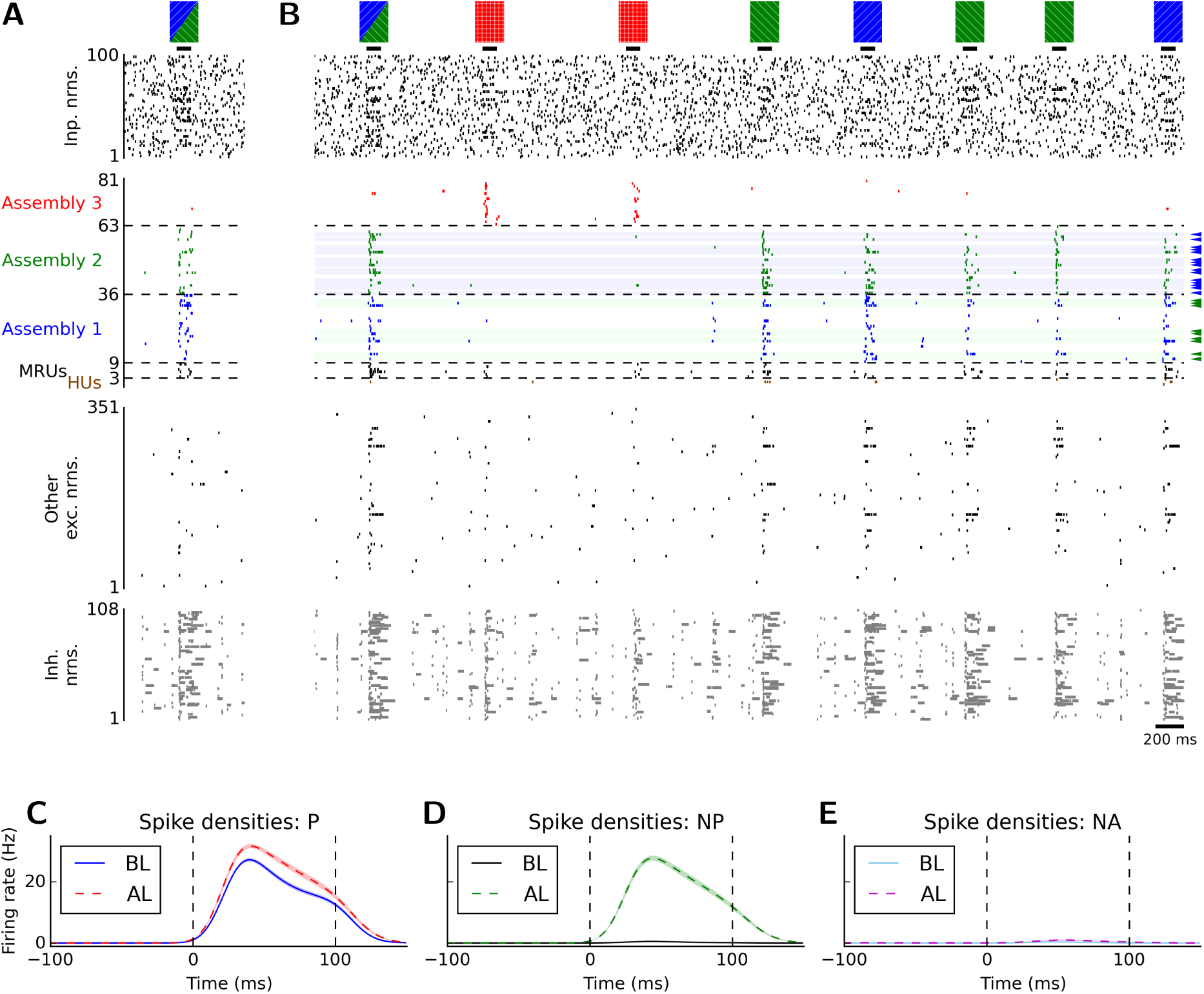
Emergence of overlap between memory traces in the model. During the association phase, the network was exposed to a combination of two previously shown rate patterns (blue and green). (A) Response of the network to the first presentation of such a combined pattern during the association phase. (B) After 20 presentations of this combined input pattern during the association phase, the green input pattern also activated a fraction of neurons from the assembly that encoded the blue input pattern, and vice versa. Such pair-coding units (PCUs; Ison et al. 2015) are indicated by a shaded background and small arrows on the right side. (C-E) Mean spike densities over all PCUs (estimated with a Gaussian kernel with *σ* = 10 ms) before (BL) and after (AL) learning of associations, averaged over 20 simulations with different random network realizations and input pattern triples in response to their (C) preferred (P), (D) non-preferred (NP), and (E) non-associated (NA) pattern. The stimulus onset was at *t* = 0 ms, dashed vertical lines indicate the pattern presentation period. Colored curves represent the mean and shaded areas represent the SEM over all 20 simulations (very small, hardly visible). After learning, a significantly increased firing rate in response to the NP stimulus, but not to the NA stimulus could be observed. Compare with Figure 5A-C of (Ison et al. 2015).

As a control, we also considered other values for some of the parameters and analyzed the impact of parameter variations on the emergence of memory traces and associations. Specifically, we analyzed in Figures 7 and S7 in the *Supplementary Material* how the results depended on the values of five selected parameters (see Section 2.5) which are thought to be important for the balance between excitation and inhibition in the recurrent network. We considered in these figures randomly generated networks (with newly drawn connectivity and initial parameters) and input patterns for a large range of parameter values.

**Figure 6:**
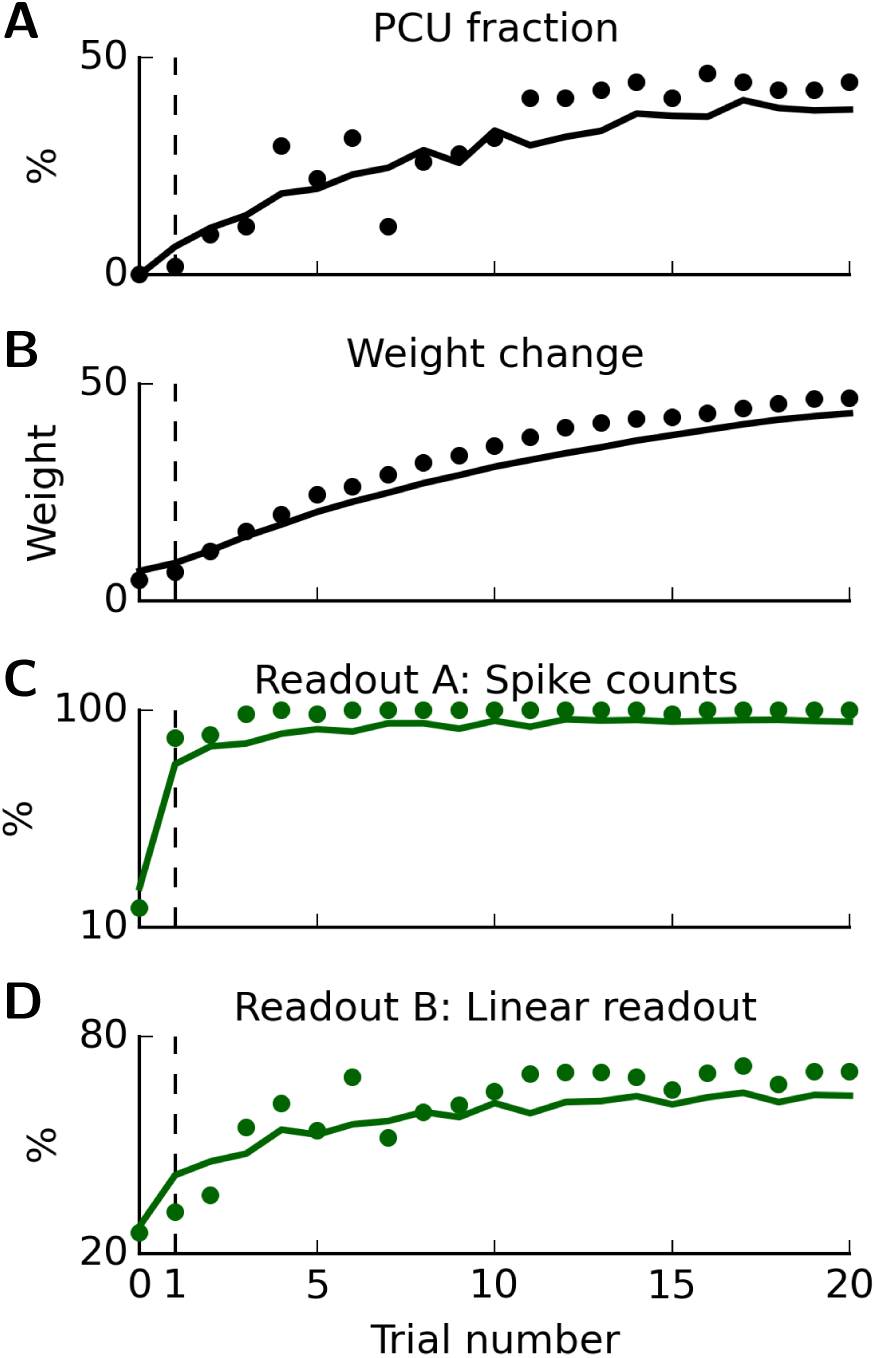
Time course of changes in synaptic weights, assembly organization, and computational function in the recurrent network during 20 presentations (“trials”) of the combined input pattern (trial 0: initial state). Solid lines represent average results over 20 simulations with different random network realizations and input pattern triples. Small circles depict results for the standard model. (A) Mean fraction of PCUs relative to their corresponding assembly sizes. (B) Mean weight changes of synaptic connections between the two associated assemblies. (C) Mean readout performance A using spike counts. (D) Mean readout performance B using a linear readout.

**Figure 7:**
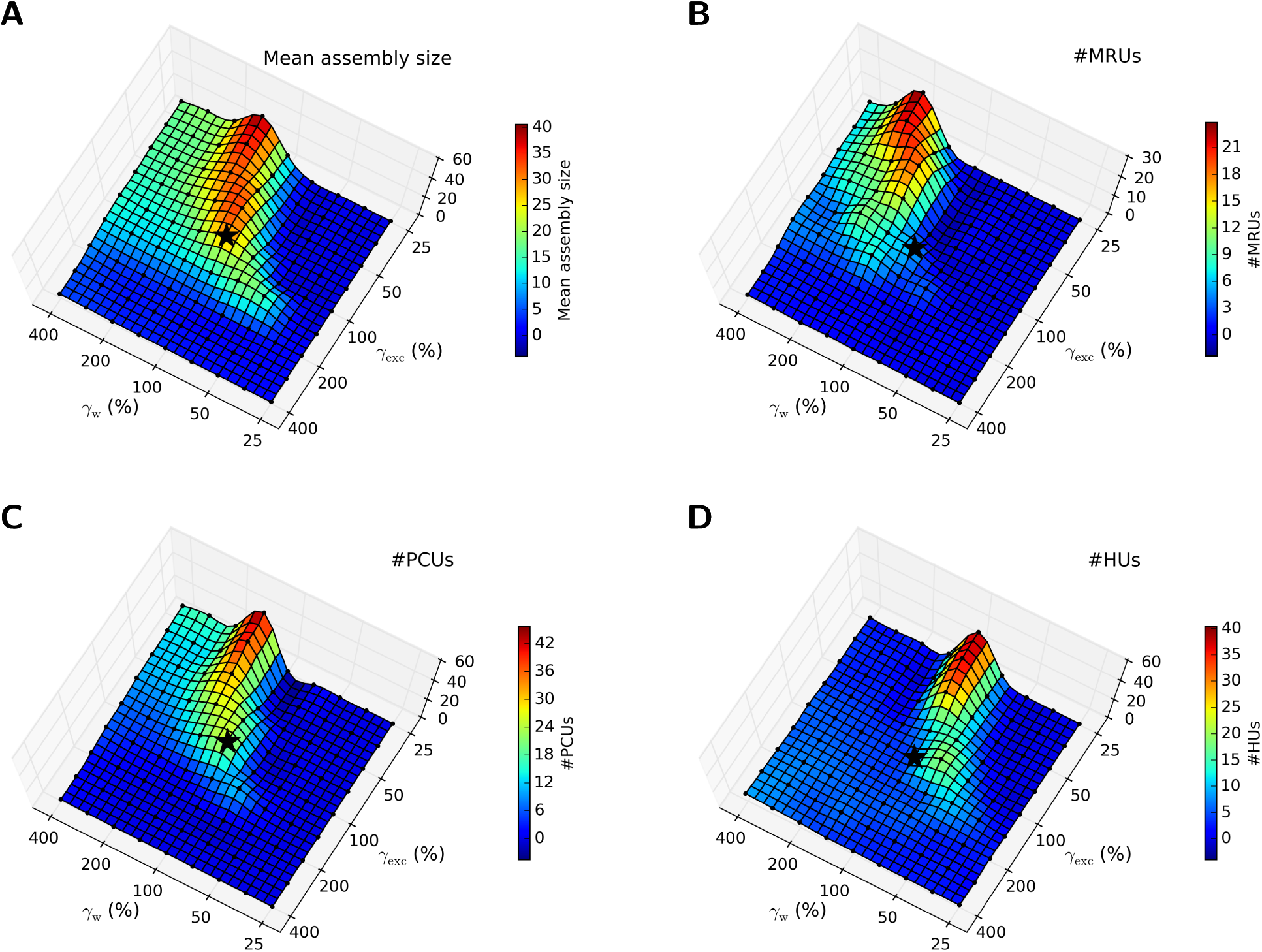
Impact of the scaling factors *γ*_exc_ and *γ*_w_ on (A) the mean assembly size, (B) the number of MRUs, (C) the number of PCUs, and (D) the number of HUs. Mean values were estimated over 10 simulations with different random seeds. The * symbols mark the values in the standard model.

#### 2.1.1 Neuron model

Our network model was a randomly connected recurrent neural network consisting of 432 excitatory and 108 inhibitory point process neurons (without adaptive threshold) with a stochastic firing criterion, to capture the natural variability of neurons. The parameters of this model were set to the mean values of the experimental data given in (Jolivet et al. 2006). In this model, the response kernel defines the shape of postsynaptic potentials (PSPs). We defined the response kernel of both excitatory and inhibitory neurons as a double-exponential function

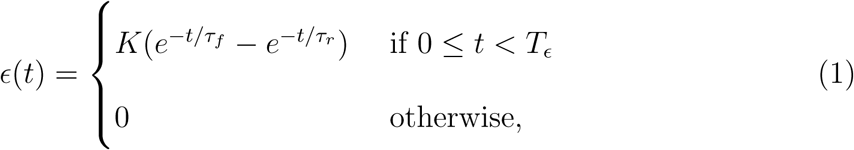

with a rise time constant of *τ*_*r*_ = 2 ms, a fall time constant of *τ*_*f*_ = 20 ms, and a cut-off at *T*_*E*_ = 100 ms. The scaling factor *K* was computed to obtain a peak value of 1. The instantaneous firing rate *r* of a neuron *i* depended on the current membrane potential *u*_*i*_(*t*) and was given in our simulations by the transfer function

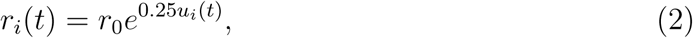

with *r*_0_ = 1.238 Hz. The membrane potential of an excitatory neuron *i* was given by

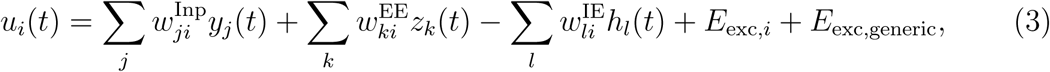

which was the weighted sum of PSPs *y*_*j*_(*t*) caused by external inputs, PSPs *z*_*k*_(*t*) caused by excitatory neurons in the recurrent circuit, PSPs *h*_*l*_(*t*) caused by inhibitory neurons, and its excitability. The term 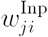 denotes synaptic weights from all input neurons *j* projecting to neuron *i*, 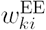 denotes recurrent synaptic weights from all excitatory neurons *k* projecting to neuron *i*, and 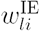 denotes synaptic weights from all inhibitory neurons *l* projecting to neuron *i* respectively. The (unweighted) sum of PSPs caused by input neuron *j* was defined as

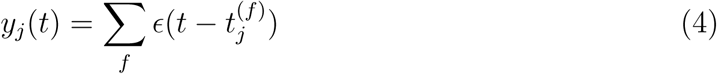

for *N* input spikes at spike times 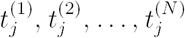. The (unweighted) sums of PSPs *z*_*k*_(*t*) and *h*_*l*_(*t*) were defined in an analogous manner. The neuronal excitability consisted of an individual component *E*_exc,*i*_ for neuron *i* drawn from a log-normal distribution across neurons with *µ* = 2.64 and *σ* = 0.23 (of the underlying normal distribution) which was shifted by *-*600 towards negative values, and a generic excitability of the excitatory population given by

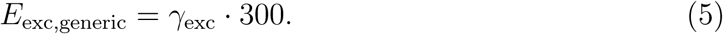

A positive scaling factor *γ*_exc_ was introduced to scale the relative contribution of the generic neuronal excitability of the excitatory population, to investigate the functional impact of certain network parameters (see Section 3.5). In the standard model, this factor was set to *γ*_exc_ = 1. The membrane potential for inhibitory neurons was defined in an analog way but without external inputs and a larger value for the generic excitability *I*_exc,generic_ = 450 of the inhibitory population. After generating a spike, neurons entered a refractory period of random length drawn from gamma distributions (shape parameter *k* = 2) with a mean value of 10 ms for excitatory and 3 ms for inhibitory neurons.

Since the voltage values in our model are unit-free (Equations (2) and (3)), we employed a heuristic approach to estimate how the voltage values in our model *u*_*model*_ would translate to biological voltages *u*_*biol*_. For this purpose, we recorded the membrane potentials of the excitatory neurons with a time resolution of 1 ms. We found a mean voltage value of 30.12 one time step before a spike was generated (this is the closest estimate, since we cannot record membrane potentials directly at the time of a spike –it is reset to zero whenever a spike is generated). Similarly, we estimated the mean baseline membrane potantial at rest of a neuron to be −905.90. Assuming a firing threshold of a biological neuron at 20 mV and a resting potential at 0 mV, this gives us an estimate of the membrane voltage *u*_*biol*_ in mV of

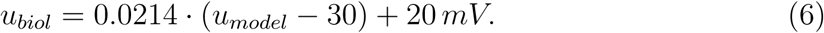

Hence one voltage unit in our model amounts to 0.0214 mV.

#### 2.1.2 Network

Figure 1A shows a schematic representation of the network architecture. A pool of 432 excitatory and a pool of 108 inhibitory neurons were reciprocally connected, while each pool was also recurrently connected. Additionally, a pool of 200 external neurons (termed input neurons) was connected to the excitatory population.

All 540 network neurons were arranged in a 3D grid of 6 *×* 6 *×* 15 neurons. The recurrent connectivity among excitatory neurons was uniform, in accordance with experimental data on the anatomy of CA3 which show a rather uniform recurrent connectivity among excitatory neurons within this area (Guzman et al. 2016). We chose in our standard model a connection probability of *p*_E*→*E_ = 50 % to assure a sufficient number of connections between excitatory neurons despite the small network size. All other connection probabilities between excitatory and inhibitory neurons were exponentially distance-dependent, resulting in very strong and local inhibition. The distance-dependent connection probabilities between pairs of neurons were defined as

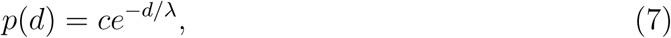

where *d* denotes the distance between the neurons in grid units, *λ* = 0.25 denotes the length scale, and *c* is a scaling parameter with values that depended on the connection type as shown in Table 1. This resulted in average connection probabilities of around 4 % for E*→*I and I*→*I, and around 5 % for I*→*E connections (100 % in close vicinity). The pool of input neurons was randomly connected to the excitatory population with a uniform connection probability of 50 %.

**Table 1:** Connection parameters of the standard model between input (Inp), excitatory (E), and inhibitory (I) pools of neurons. The connection probabilities were either constant (uniform) or exponentially distance-dependent with parameters *c* and *λ*. Initial weights were drawn from gamma distributions, synaptic delays from normal distributions with given means *µ* and CVs and a positive scaling factor *γ*_w_ = 1.

| Connection | Conn. prob. | Dist.-dep. | Init. weights | Delays |
| --- | --- | --- | --- | --- |
| Inp→E | 50 % | *uniform* | $\mu = \gamma_w \cdot 15$<br>$CV = 0.7$ | $\mu = 5 \text{ ms}$<br>$CV = 0.5$ |
| E→E | 50 % | *uniform* | $\mu = \gamma_w \cdot 2.5$<br>$CV = 0.7$ | $\mu = 5 \text{ ms}$<br>$CV = 0.5$ |
| E→I | 4 % | $c = 2 \cdot 10^5$<br>$\lambda = 0.25$ | $\mu = 1000$<br>$CV = 0.7$ | $\mu = 2 \text{ ms}$<br>$CV = 0.5$ |
| I→E | 5 % | $c = 4 \cdot 10^5$<br>$\lambda = 0.25$ | $\mu = 1375$<br>$CV = 0.7$ | $\mu = 2 \text{ ms}$<br>$CV = 0.5$ |
| I→I | 4 % | $c = 1 \cdot 10^5$<br>$\lambda = 0.25$ | $\mu = 6000$<br>$CV = 0.7$ | $\mu = 2 \text{ ms}$<br>$CV = 0.5$ |

**Table 2:**
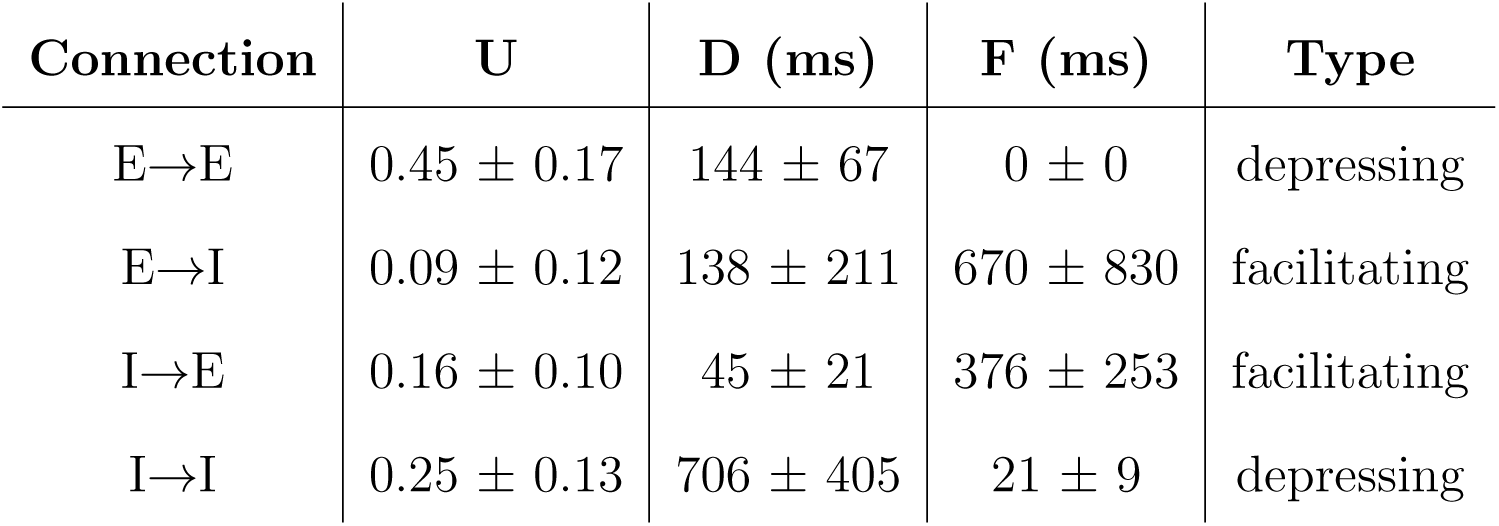
Parameters for short-term plasticity in the model: U (release probability), D (time constant for depression), and F (time constant for facilitation). Mean ± SD values were taken from experimental results given in (Testa-Silva et al. 2014; E*→*E connections), (Gupta et al. 2000; I*→*E connections) and (Markram et al. 2015; remaining connections) respectively.

Synaptic delays for input and recurrent excitatory connections, as well as delays to and from inhibitory neurons were drawn from normal distributions with mean values as given in Table 1 and a coefficient of variation *CV* = 0.5.

Initial synaptic weights were drawn from gamma distributions with mean values as given in Table 1 and *CV* = 0.7. Specifically, the mean initial input weights 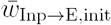 and the mean initial recurrent excitatory weights 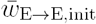 were given by

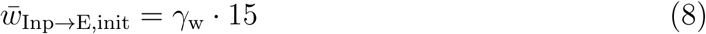

and

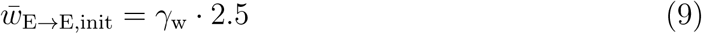

respectively. A positive scaling factor *γ*_w_ was introduced to scale the contributions of the synaptic weights between input and excitatory as well as between excitatory neurons relative to the other weights in the network, to investigate the functional impact of certain network parameters (see Section 3.5). In the standard model, this factor was set to *γ*_w_ = 1. From the modelling perspective it was important for the generation of memory traces (see Section 3.2) to use initial recurrent weights lower than input weights so that the network operated in a stable input-driven regime from the beginning, and to allow recurrent weight to successively increase through long-term plasticity processes (see Section 2.1.4). After initialization, all weights between excitatory and inhibitory neurons in the circuit were adjusted as described in Section 2.1.3 to account for effects of short-term synaptic plasticity.

#### 2.1.3 Short-term synaptic plasticity

Our model for synaptic connections included data-constrained short-term plasticity, i.e., a mixture of paired-pulse depression and facilitation that depended on the type of the pre- and postsynaptic neuron. This can be described by three parameters (Markram et al. 1998): U (release probability), D (time constant for depression), and F (time constant for facilitation). The PSP generated by the n-th presynaptic spike in a spike train with a time interval Δ*t* between the (n-1)-th and n-th spike is given by

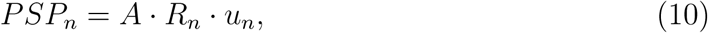

where *A* is the absolute synaptic efficacy and *R*_*n*_ and *u*_*n*_ are defined by

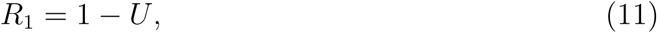

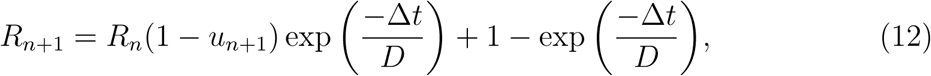

and

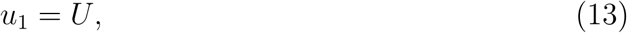

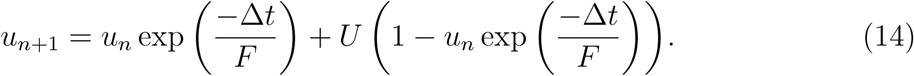

Values of UDF parameters for synapses between different cell types and layers in the somatosensory cortex of rats have been reported in (Gupta et al. 2000) and more recently in (Markram et al. 2015). Additionally, for the adult human brain (Testa-Silva et al. 2014) find in synaptic connections between layer 2/3 pyramidal neurons frequency-dependent depression but no facilitation. In our model, values for UDF parameters were drawn from bounded gamma distributions with mean values and standard deviations (SD) for E*→*E synapses taken from human experimental data (Testa-Silva et al. 2014), for E*→*I and I*→*I synapses taken from the range of values among the most frequent connection types of the recent experimental results reported in (Markram et al. 2015), and for I*→*E synapses taken from (Gupta et al. 2000). The UDF parameters were bound between [0.001, 0.999] (parameter U) and [0.1 ms, 5000 ms] (parameters D, F) respectively. The used mean and SD values are summarized in Table 2 and the corresponding distributions are illustrated in the three-dimensional UDF parameter space in Figure 1B.

After random initialization of the weights and UDF parameters according to the distributions specified above, all weights were adjusted based on their steady-state values in the following way (Markram et al. 1998, Sussillo et al. 2007). For a given constant presynaptic firing rate *f*_0_, steady-state values of the synaptic weights 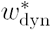 of the dynamic synapses can be computed as

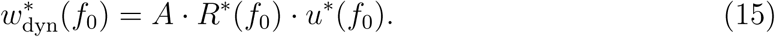

The steady-state values for the synaptic availability *R* and the synaptic utilization *u* at a given rate *r* are given by

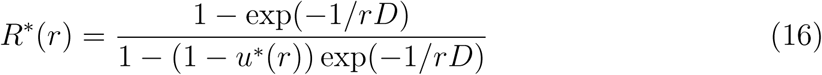

and

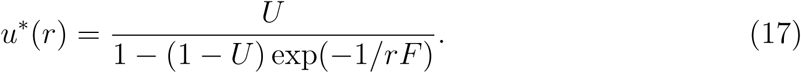

Based on these steady-state values, the initial weight *w*_init_ (absolute synaptic efficacy) of each dynamic synapse in the network was adjusted so that its dynamic weight at an assumed constant presynaptic firing rate *f*_0_ = 5 Hz (which is the noise rate of the input neurons when no input patterns are presented; see Section 2.2.1) corresponded to its previously assigned initial weight value, by

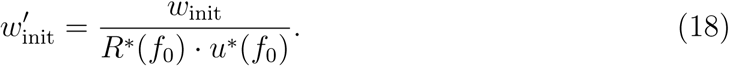

This adjustment of initial weights was beneficial to reduce initial transients and to decrease the burn-in time of the network needed to reach steady-state activity after starting the simulation.

#### 2.1.4 Long-term synaptic plasticity

Synaptic connections from input to excitatory as well as between excitatory neurons in our model were subject to a data-constrained rule for STDP: the triplet rule (Pfister and Gerstner 2006). This rule was implemented through the `stdp triplet synapse` model provided in NEST, implementing all-to-all interactions (Pfister and Gerstner 2006). Weight updates for each pre- and postsynaptic spike arriving at time *t*^pre^ and *t*^post^ respectively were given by

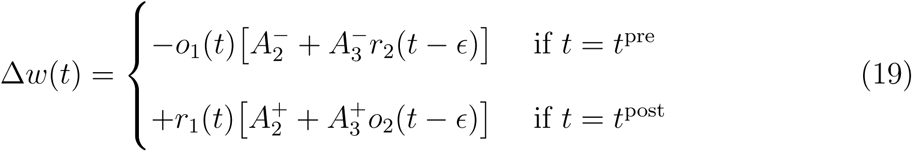

The variables *r*_1_ and *r*_2_ were presynaptic and *o*_1_ and *o*_2_ were postsynaptic detector variables which were increased by 1 upon pre- and postsynaptic spike arrival and decayed with time constants 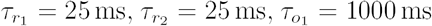 and 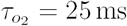 respectively. The amplitude parameters were chosen as 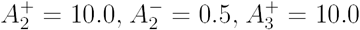 and 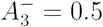 respectively. The symbol *E* denotes a small positive constant indicating that weight updates were done before updating the detectors *r*_2_ and *o*_2_. The amplitude parameters and time constants were qualitatively chosen based on the experimental data in (Pfister and Gerstner 2006; hippocampal culture set) but then adapted to the magnitude of weights and temporal dynamics of our model so that the emergence of stable memory traces through STDP (see Section 3.2) for the given input patterns was possible. The STDP induction rule is illustrated in Figure 1C in a typical pairing experiment with a pairing frequency of 20 Hz. Synapses from input neurons to excitatory neurons as well as between excitatory neurons in the network were subject to the same plasticity rule. Total relative weight changes were limited to [0 %, 200 %] for input synapses and to [0 %, 1000 %] for recurrent excitatory synapses respectively with respect to weight initialization.

### 2.2 Details to emergent memory traces

#### 2.2.1 Input patterns

In each simulation, we repeatedly presented three input patterns to the network (see Section 2.2.3). In total, we generated 20 different random network realizations each of which had a specific triple of input patterns. Each input pattern was a sparse stationary rate pattern over 100 ms with 20 of 200 randomly drawn on-channels (i.e., 10 % of all channels) with firing rates of *r*_on_ = 40 Hz, while all other 180 off-channels remained silent (i.e., *r*_off_ = 0 Hz). On-channels were drawn in such a way that the maximum overlap between patterns (i.e., number of shared channels) in a triple was limited to 2 (i.e., 10 % of on-channels).

From each pattern triple, we chose an arbitrary pattern and termed it the *blue* pattern, another one the *green* pattern, and the third one the *red* pattern for easy reference in the text. As detailed below, a neural assembly emerged for each of these patterns due to synaptic plasticity, and these were termed the *blue*, the *green*, and the *red* assembly respectively.

At each presentation of such a pattern, fresh Poisson spike trains were independently generated for each of its on-channels with a rate of *r*_on_. In addition, each of the 200 input channels was superimposed with a freshly generated Poisson spike train with a rate of 3 Hz. In periods where no input patterns were presented, input neurons emitted freshly generated Poisson spike trains at 5 Hz.

#### 2.2.2 Network simulation and analysis

All network simulations and data analyses were done in Python 2.7.13 using the NEST Simulator 2.12.0 (Kunkel et al. 2017) with some user-defined optimizations, together with the PyNEST Interface (Eppler et al. 2009). A time resolution of 1 ms was used for all simulations.

The experimental setup consisted of the main simulation and various analyses of the network, as depicted in Figure 2 and described in the following.

##### Main simulation

The main simulation consisted of five input phases during which STDP was always active for synapses from input to excitatory as well as between excitatory neurons. The different phases differed only in their length and in the type of input patterns that were presented to the network. An overview of the sequence of used input phases is given below, more details can be found in the respective sections as indicated:

1. Initialization phase (250 s), during which only noise was presented (no patterns).
2. Encoding phase for the emergence of memory traces (250 s), during which three input patterns were randomly presented to the network (see Section 2.2).
3. Resting phase 1 (250 s), during which only noise was presented.
4. Association phase for the emergence of associations (36 s), during which a combined pattern (blue and green) was repeatedly presented to the network (see Section 2.3).
5. Resting phase 2 (250 s), during which only noise was presented.

##### Network analyses

Different aspects of the network at certain time points during the main simulation (as indicated in Figure 2) were analyzed (or just plotted) in separate simulation runs. For this purpose, synaptic weights were kept constant (i.e., STDP disabled) while different types of input patterns were applied to the network. In all analyses, the network was generated in exactly the same way as in the main simulation using the same global random seed for the connection matrix, the initial parameter values, and the specific realization of the input patterns. After generating the network, new initial weights were loaded which had been extracted at a certain time point from the main simulation. So the initial state of the network corresponded to the state of the network during the main simulation at the respective time point the weights had been extracted from. Only internal state variables of the network, e.g. regarding short-term plasticity, were not identical to the main simulation. In addition, the sequence of presented input patterns differed from the main simulation by choosing another random seed for the pattern sequence generation, but which was the same for all analyses. Specifically, separate network analyses without STDP were conducted for illustration purposes of the network activity shown in Figures 3, 5B-E, and S6B. More details about what other types of analyses were conducted in this way can be found in the following sections.

#### 2.2.3 Encoding phase for the emergence of memory traces

During the encoding phase for the emergence of memory traces –which lasted for around 250 s– input patterns were presented at random time points to the network as input. Each pattern lasted for 100 ms. After a pattern presentation, a time period was randomly chosen from the uniform distribution between 0.5 s and 3 s during which input neurons emitted freshly generated Poisson spike trains at 5 Hz. After this noise period, the next pattern was randomly chosen and presented and so on. The first pattern at the beginning of a phase was chosen randomly with a uniform probability over the three patterns (but not presented to the network; see below). Subsequent patterns were drawn based on a switching probability of 75 %, meaning that with probability 75 %, there was a switch to another pattern (drawn uniformly from the other patterns), and with 25 % probability the same pattern was repeated.

When generating such a sequence of input patterns and noise periods of a given length, the exact duration of the resulting input phase was determined by the following rules: The first pattern was always omitted so that each sequence started with a noise period at the beginning. In case there was a pattern exactly at the end of a sequence, also the last pattern was omitted so that each sequence ended with a noise period. If there was already a noise period at the end of a sequence, this noise period was not truncated. So when referring to an input phase of 250 s, the exact duration could be (with a pattern duration of 100 ms) in a range between [249.8 s, 253.0 s] and was 250.85 s on average. The same rules apply to all other input phases involving input patterns accordingly.

#### 2.2.4 Definition of assembly neurons

The experiments of (Ison et al. 2015) studied the formation of associations between memory traces for a specific set of images. Neurons were classified as belonging to the memory trace (assembly) for a specific image if they significantly responded to this image based on a Wilcoxon rank-sum test (*p <* 0.05) between baseline and response intervals. As additional criterion, a median firing rate in response intervals across trials of at least 2 Hz was required. Neurons that satisfied these criteria for any of the images were termed visually responsive units (VRUs).

In an analogous manner we defined that a neuron in our recurrent network belonged to the assembly for a particular input pattern if it satisfied the same firing rate criterion and significantly responded to this input pattern based on the same Wilcoxon rank-sum test. For this purpose, the state of the network after the encoding and subsequent resting phase 1 (i.e., directly before the association phase) was analyzed (see Section 2.2.2) to identify assembly neurons. We ran this network analysis for around 325 s (see exact rules in Section 2.2.3) during which the blue, green, red, and the combined pattern (blue and green) were repeatedly presented to the network. We extracted the average firing rates for each excitatory neuron within the baseline interval [-100 ms, 0 ms] and response interval [10 ms, 110 ms] relative to the onsets of each blue, green, and red pattern presentation (termed trial; between 33 and 44 trials per pattern). Based on this, we computed the median firing rate in the response interval across all trials. Using a Wilcoxon rank-sum test (*p <* 0.05) we tested if the firing rate of a neuron across all trials was significantly higher in the response than in the baseline interval for the blue, green, and red input patterns. A neuron that showed a significant response to an input pattern and had a median firing rate in the response interval of at least 2 Hz was defined as an assembly neuron for the present input pattern, called the preferred pattern.

We refer to the neurons that belonged to the assembly for at least one of the three input patterns as pattern responsive units (PRUs). Neurons belonging to more than one assembly are referred to as multi-responsive units (MRUs).

### 2.3 Details to emergent associations

#### 2.3.1 Association phase for the emergence of associations

During the association phase, a combined pattern was repeatedly presented to the network. The sequence of input patterns and noise periods was randomly generated according to the same rules as described in Section 2.2.3 for a duration of around 36 s, but using only the combined input pattern. However, in contrast to all other phases, exactly 20 patterns were presented. Since the whole sequence of input patterns and noise periods was precomputed beforehand at the beginning of the simulation, this could be done by repeating the random generation process until the number of input patterns was 20, resulting in an average duration of 36.97 s.

Here, we always combined the blue and the green pattern (the identity of which was arbitrarily assigned before as discussed above). The combined pattern was constructed by adding the stationary rates of the 200 external input neurons for the two input patterns. For the overlapping channels between the two patterns (0-2 channels), the resulting firing rates were truncated again at 40 Hz so that all channels in the combined pattern were either on-channels with *r*_on_ = 40 Hz (38-40 channels, consisting of the on-channels of the blue and green pattern) or off-channels with *r*_off_ = 0 Hz (160-162 channels). Again, during a presentation of a combined pattern, fresh Poisson spike trains were independently generated for each of its on-channels with a rate of *r*_on_, while each of the 200 input channels was superimposed with a freshly generated Poisson spike train at 3 Hz.

#### 2.3.2 Definition of pair-coding units (PCUs)

PCUs were determined in a similar way as assembly neurons by analyzing the network directly after the association phase (see Section 2.2.2). We ran this network analysis for around 325 s (see exact rules in Section 2.2.3) with the blue, green, red, and the combined pattern (blue and green) as used in the association phase (see Section 2.3.1) repeadetly presented to the network. The average firing rates for each excitatory neuron within the baseline and response intervals (as defined in Section 2.2.4) relative to the onsets of the blue, green, and red pattern presentations were extracted (between 33 and 44 trials per pattern). Again, we used a Wilcoxon rank-sum test (*p <* 0.05) to test if the firing rate of a neuron across all trials was significantly higher in the response than in the baseline interval for the blue, green, and red input patterns.

We only considered PRUs that had exactly one of the components of the combined pattern as their preferred (P) stimulus, i.e., the blue (green) input pattern for neurons belonging to the blue (green) assembly. Accordingly, the other component was defined as their non-preferred (NP) stimulus, i.e., the green (blue) input pattern for neurons belonging to the blue (green) assembly. The red input pattern (which was not part of the combined pattern) was defined as the non-associated (NA) stimulus for this subset of PRUs.

PCUs were defined as PRUs within this subset that had a non-significant response before and a significant response after the association phase to their NP stimulus. Additionally, single-trial increases after the association phase in the response intervals of the NP stimulus were required to be significantly larger (Wilcoxon rank-sum test; *p <* 0.05) than the ones of the NA stimulus. Single-trial increases were computed as the firing rates in the response interval of a given stimulus during the network analysis after minus the mean firing rates in the response interval of this stimulus over all trials during the network analysis before the association phase. So PCU were identified in the same way as described in the experimental procedures in (Ison et al. 2015), but with different baseline and response intervals. Instead of baseline and response intervals of [-500 ms, 100 ms] and [200 ms, 800 ms] respectively relative to stimulus onsets in (Ison et al. 2015) we chose [-100 ms, 0 ms] and [10 ms, 110 ms] respectively. This was done to account for the shorter pattern presentation interval of 100 ms in our model as compared to single pictures shown for 1000 ms to the participants in (Ison et al. 2015). As an additional constraint we excluded neurons which showed no longer a significant response to the P stimulus after the association phase (*<* 1 neuron on average out of all PCUs identified otherwise).

#### 2.3.3 Stability of the emergent number of PCUs

Over 20 simulations with different network initializations and input patterns, we found an average number of 22.2 ± 7.0 SD (min: 11; max: 35) PCUs. We found that after the formation of associations, 18.1 ± 5.3 SD previously non-responsive units (to any of the three input patterns) became responsive to a single input pattern and 5.7 ± 2.5 SD units, so-called hierarchical units (HUs; exact definition in Section 2.3.4), to the combined pattern respectively. Over all 20 simulations we found a weak but significant linear correlation between the assembly sizes and the resulting numbers of PCUs per assembly (*r* = 0.45, *p <* 0.01). Details can be found in Figures S3 and S4 in the *Supplementary Material*.

#### 2.3.4 Definition of hierarchical units (HUs)

In hierarchical memory models (Kahana et al. 2008, Kahana 2012), a new memory trace is thought to emerge for the combined input pattern in form of HUs responding only to the combined pattern. We tested whether such HUs existed and defined them as units that did not respond to any separate pattern before the formation of associations (i.e., non-PRUs), but became responsive to the combined pattern only (but not to any separate pattern) after the emergence of associations. Responsiveness to the combined pattern was determined in the same network analysis that was conducted to identify PCUs (during which also combined patterns were presented; see Section 2.3.2)) using the same criteria as for assembly neurons defined in Section 2.2.4. According to this definition, HUs also included rare cases of units that were responsive to the combined pattern even before the association phase (*<* 1 neuron on average).

### 2.4 Details to neuronal and functional learning curves

Behavioral performance of learned associations in (Ison et al. 2015) was tested during learning by showing an image and asking the participant to select the corresponding associated image from a list of images (Task 3). Similarly, in our network model, we investigated after which combined pattern presentation in the association phase (see Section 2.2.2) the formed associations became functionally useful, in the sense that a downstream network could infer the associated pattern for a given input pattern. For this purpose, we ran stepwise network analyses after an increasing number of combined input patterns (from 0-20) presented during the association phase. Weights were extracted from the main simulation at intermediate time points directly before the pattern onsets of all 20 combined patterns as well as at the end of the association phase, to be used in these network analyses. We then extracted the resulting numbers of PCUs, weight changes between the associated assemblies, and readout performance using two different kinds of readout strategies.

#### 2.4.1 Readout A: Readout based-on spike counts

This readout can be seen as mimicking the task to select the associated image from a list of images in a multiple choice test in the experiments of (Ison et al. 2015). Spike counts from all neurons belonging to an assembly were extracted from the baseline interval of [-100 ms, 0 ms] and the response interval of [10 ms, 110 ms] relative to stimulus onsets of the blue, green, and red patterns. These spike counts were determined in all of the stepwise network analyses after the presentation of increasing numbers of combined input patterns during the association phase (between 33 and 44 trials per pattern in each analysis). For each assembly, the mean spike count differences between the response and the baseline interval over all neurons belonging to the corresponding assembly were computed. When presenting a blue (green) input pattern, the spike count differences for its associated (green (blue)) and non-associated (red) assembly were compared, and the assembly with a higher spike count difference was selected as output of the readout. The total readout performance was estimated by computing the fraction of times (of all blue (green) trials) the correct (i.e., the associated) assembly was selected.

#### 2.4.2 Readout B: Linear readout

For each of the three assemblies (red, blue, and green), a linear readout was trained with the standard algorithm for a support vector machine with parameter *C* = 1 to detect its activation in response to its preferred (P) input pattern. We used the LinearSVC implementation from the scikit-learn 0.18 library (Pedregosa et al. 2011). 432-dimensional feature vectors for this readout were extracted by taking the non-weighted PSPs (similar to *z*_*k*_(*t*) in Equation (3), but with longer time constants *τ*_*r*_ = 4 ms, *τ*_*f*_ = 40 ms, and *T*_*c*_ = 200 ms, to get a more stable output of the readout) summed over time at time point 100 ms relative to pattern onsets from all excitatory neurons (i.e., without any prior knowledge about the actual assembly neurons). Each classifier was then trained on 50 % of the data to distinguish target patterns (i.e., assembly activation at its preferred pattern presentation; around 19 presentations) from non-target patterns (around 39 presentations) during the network analysis directly before the association phase (i.e., the same analysis assembly neurons were determined in; see Section 2.2.4).

We then tested whether this readout was able to detect during the network analysis directly after the association phase (i.e., the same analysis PCUs were determined in; see Section 2.3.2) an indirect activation of its corresponding assembly via its associated non-preferred input pattern (again, based on around 19 target and 39 non-target pattern presentations). To take unbalanced classes into account (there were always twice as many non-target as target patterns), the class weight of the smaller class was increased by a factor of 2, using the `class weight` argument of LinearSVC. Details of the LinearSVC implementation can be found in the web-based API documentation of scikit-learn (LinearSVC 2016). We used the *balanced accuracy* as performance measure which is defined as 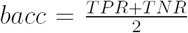, with TPR being the true positive rate and TNR the true negative rate. The TPR (TNR) is defined as the fraction between the number of samples that were correctly classified as positive (negative) examples and the total number of positive (negative) examples in the test set. Moreover, to increase the sensitivity of the readouts to detecting associations, the separating hyperplane was shifted towards the non-target class by decreasing the corresponding class weight by a factor of 1 *·* 10^*-*3^.

### 2.5 Details to the functional impact of network parameters

In Section 2.1, we described the network parameters and the chosen standard values we used throughout our simulations. We investigated the impact of certain network parameters and the robustness of the results against variations of network initialization (i.e., specific network connectivity and initial parameter values) and input patterns. To this end, we ran extensive computer simulations and varied one or two parameters at a time while keeping the standard values for all other parameters. To have comparable ranges of values despite the fact that different parameters have different scales and units, we used relative parameter changes with respect to the corresponding standard value. To cover a reasonably wide range of values, we used nine logarithmically spaced steps from 25 % to 400 % of the standard parameter values. When two parameters at a time were varied, 9 *×* 9 grid points were computed and intermediate values were estimated by bicubic interpolation to create 2D surface plots. For the connection probability between excitatory neurons we used absolute probability values in nine linearly spaced steps from 10 % to 90 %.

All simulations were done using 10 different global random seeds, having an effect on both network generation (e.g., connection matrix, initial parameter values) and the specific realization of input patterns. The average results over these 10 simulations are reported. Parameters that were investigated are summarized in Table 3, together with short descriptions and their standard values, and include *E*_exc,generic_, *I*_exc,generic_, *p*_E*→*E_, *γ*_exc_, and *γ*_w_. In a subset of the simulations, the combined scaling factor *γ* := *γ*_exc_ = *γ*_w_, which was not an intrinsic parameter of the network, was treated as single parameter used to simultaneously control both scaling factors, *γ*_exc_ and *γ*_w_, in the same way. Results for variations of parameters *γ*_exc_ and *γ*_w_ can be found in Figure 7 while results for parameters *E*_exc,generic_, *I*_exc,generic_, *γ*_w_, and *p*_E*→*E_ can be found in Figure S7 in the *Supplementary Material*.

**Table 3:** List of selected network parameters that were varied to investigate their functional impact, with a short description and their standard values. In a subset of simulations, the combined scaling factor *γ* was treated as single parameter used to simultaneously control the scaling factors *γ*_exc_ and *γ*_w_.

| Symbol | Parameter description | Standard value |
| --- | --- | --- |
| $E_{\text{exc,generic}}$ | Generic excitability of the excitatory population | 300 |
| $I_{\text{exc,generic}}$ | Generic excitability of the inhibitory population | 450 |
| $p_{\text{E} \rightarrow \text{E}}$ | E→E connection probability | 50 % |
| $\gamma_{\text{exc}}$ | Scaling factor of generic neuronal excitability of the excitatory population | 1.0 |
| $\gamma_{\text{w}}$ | Scaling factor of initial synaptic weights between Inp→E and E→E connections | 1.0 |
| -----<br>$\gamma$ | Combined scaling factor $\gamma := \gamma_{\text{exc}} = \gamma_{\text{w}}$ | -----<br>1.0 |

### 2.6 Details to the balance between excitation and inhibition

The E/I balance based on membrane potentials was continuously measured during the main simulation. Following the general definition of Okun and Lampl (2009), we computed the E/I balance as the mean ratio over all excitatory neurons of the excitatory (i.e., positive) and inhibitory (i.e., negative) contributions to the membrane potentials *u*_*i*_ (see Equation (3)), which is arguably the most direct way of measuring the relative contributions of excitatory and inhibitory synaptic inputs to a neuron. Since the total neuronal excitabilities *E*_exc,total,*i*_ = *E*_exc,*i*_ + *E*_exc,generic_ across neurons *i* were usually negative, they were counted as inhibitory contribution. In case values of *E*_exc,total,*i*_ exceeded zero (when testing the functional impact of network parameters; see Section 2.5) they were counted as excitatory contribution. For this purpose, the total excitability of neuron *i* was splitted into its positive and negative contributions, using

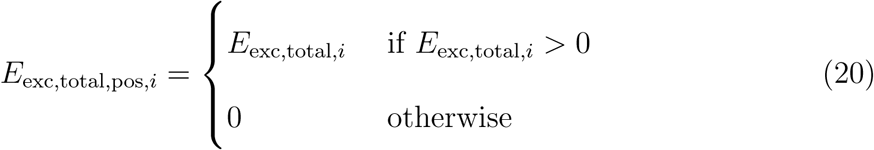

and

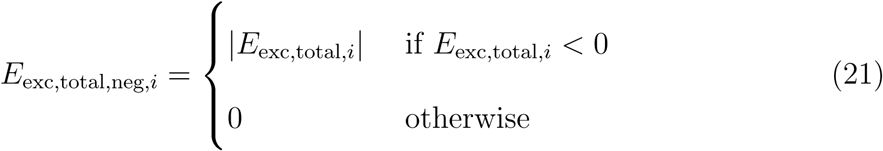

respectively. To avoid numerical problems (divisions by zero), we first divided the excitatory by the inhibitory contributions, since the inhibitory contributions were usually never zero due to the negative generic excitability *E*_exc,generic_ (except for at spike times where the membrane potential was reset to 0 mV for one time step). We then averaged over all excitatory neurons (excluding neurons where this ratio was not finite in a given time step) and smoothed the resulting mean ratio over time with a moving average (MAV) filter with a filter length of 10 s to get a stable continuous estimate. Finally, since inhibition was larger than excitation, we inverted this smoothed curve, resulting in an I-to-E ratio of

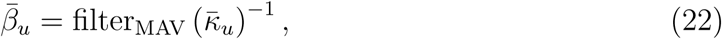

with

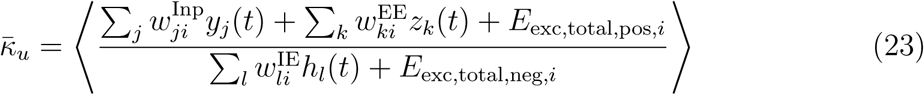

Discrete values of 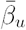 were extracted from this inverted smoothed curve at three points of interest during the course of the simulation (compensated for filter delays): (i) after the initialization phase, (ii) directly before, and (iii) directly after the association phase (see Section 2.2.2).

## 3. Results

### 3.1 The network model

We examine emergent neural codes for the association of memory items in a generic recurrent network of excitatory and inhibitory spiking neurons. In contrast to artificial neural network models, one can readily compare network responses and neural codes of such a model with experimental data from electrode recordings in the brain, since both have the form of spike trains and spike rasters. Similarly to the model for area CA3 of (Guzman et al. 2016) we assume that the connection probability between excitatory neurons is distance-independent. However the probabilities of all other types of synaptic connections were assumed to drop exponentially with the distance between the somata of neurons, which were arranged on a 3D grid. We included data-constrained short-term plasticity of synapses in the model because it has a significant impact on the network dynamics. In particular, the short-term plasticity of synaptic connections between excitatory neurons was modelled according to data from the human brain (Testa-Silva et al. 2014), which showed that these synapses are depressing. In addition, we used a standard data-constrained rule for long-term spike-timing-dependent plasticity (STDP) of synapses between excitatory neurons, the triplet rule of (Pfister and Gerstner 2006).

### 3.2 Generation of memory traces

The experiments of (Ison et al. 2015) studied fast formation of associations between two unrelated memory items, represented through images of familiar faces and landscapes. One commonly assumes that each such memory item is represented in the human MTL by a sparse distributed set of concept cells, often referred to as memory trace or assembly of neurons (Quian Quiroga 2016). In order to study the formations of associations, we first needed to generate such memory traces in our model through STDP. We reasoned that such a memory trace should emerge for each sufficiently often occurring external input to the model. This external input was produced by a separate pool of excitatory spiking neurons, that had randomly generated connections to neurons in the recurrent network.

#### Input patterns

We chose 3 firing rate patterns of 200 external Poisson neurons as input patterns, referred to as the blue, green, and red pattern. Resulting spike inputs were freshly generated for each presentation, and superimposed with freshly generated noise at 3 Hz for each input neuron, see Section 2.2.1 for details.

After an initialization phase of 250 s without an external input, each of the three input patterns was represented repeatedly and interlaced to the network, at random time points within a continuous spike input stream over 250 s (Fig. 2). Each input pattern was presented on average 39 times. During this “encoding phase”, in which the emergence of memory traces through STDP was studied, the three input patterns with superimposed noise were interleaved with periods of pure noise: freshly generated Poisson spike trains at 5 Hz, with randomly drawn durations between 0.5 s and 3 s (see the top row of Figure 3).

More precisely, we analyzed the emergence of pattern responsive units (PRUs). These were defined exactly as in (Ison et al. 2015) as neurons that significantly responded to at least one of the three input patterns. We refer to the set of neurons in the network that preferentially responded after this encoding and a subsequent resting phase 1, see Figure 2, to one of the three input patterns as the blue, green, and red assembly (see second row of Figure 3, (Future) Assembly 1-3). Figures 3A and B illustrate the network activity in response to the three input patterns (A) at the initial state of the network before any plasticity took place and (B) after the encoding and subsequent 250 s resting phase 1, which is the time point at which PRUs and assembly neurons were determined (see Section 2.2.4). Overall the network response was quite sparse, as found in most recordings from cortex, with inhibitory neurons being more active than excitatory neurons. Furthermore the network response became even sparser once memory traces were generated, see Figure 3B. A sparsening of responses is commonly reported as an impact of perceptual learning (Hoffman and Logothetis 2009, Lim et al. 2015; see also references therein). We found out through a control experiment (not shown) that the chosen primarily local connectivity of inhibitory neurons was essential for the emergence of assembly codes. For the sake of completeness we show in Figure 3 also the response of a very small number of multi-responsive units (MRUs), defined as in (Ison et al. 2015), and of hierarchical units (HUs, hardly visible in Figure 3). These populations of neurons will become relevant for the emergence of neural codes for the association of two input patters, see Sections 3.3 and 3.5.

The evolution of synaptic weights under STDP, which explains the difference between the network responses in panels A and B of Figure 3, is shown in Figure 4A and in the upper three traces of Figure 4B. Weights from input neurons to neurons in an assembly *k* (see Fig. 3A) were computed by taking only those input neurons into account which fired during pattern *k* at a high rate; *k* = 1, 2, 3. Figure 3B shows that the weights of synaptic connections within an assembly increased significantly during the encoding phase but saturate towards its end since many weights reach their maximum value. Doubling the length of the encoding phase did not lead to a significantly higher saturation level (not

The three assemblies that emerged were rather sparse and distributed over the whole 3D volume of the network, with their means approximately centered at the center of the volume. In this trial STDP gave rise to 78 PRUs, with assemblies of sizes 27, 27, 18 for the three input patterns. The exact assembly sizes and numbers of PRUs and MRUs depended on the random choice of the network initialization and input patterns (see Section 3.5). Twenty simulations with random networks realizations (with newly drawn connectivity and initial parameters) and new input patterns yielded average assembly sizes of 28.4 ± 5.6 SD (min: 17; max: 45). The total number of PRUs was found to be 91.2 ± 10.4 SD (min: 67; max: 111), the number of MRUs 6.1 ± 4.3 SD (min: 0; max: 16). Relative differences between the three assembly sizes were measured by computing their standard deviation for each simulation individually, which resulted in a mean standard deviation of 4.5 over all 20 simulations. An overview over these 20 simulations can be found in Figure S1 in the *Supplementary Material*.

The emergence of assemblies as memory traces in networks of spiking neurons has already previously been modelled (Klampfl and Maass 2013, Litwin-Kumar and Doiron 2014, Zenke et al. 2015). In comparison with these preceding models we used here a simpler model that did not require a specific connectivity structure, homeostasis, or long-term plasticity of inhibitory synapses. The neural recordings of (Ison et al. 2015) show that the firing activity of neurons that belong to an assembly tend to return to baseline activity soon after the stimulus that evoked this memory trace has been removed. This feature is duplicated in our model, see Figure 3B.

We would like to mention on the side that the network can learn to generate, with the same plasticity rule, also assemblies for new input patterns that are introduced after the encoding phase, see Figure S2A in the *Supplementary Material*. This turned out to have little effect on the three previously generated assemblies (see Figure S2A and B). Through another control experiment (not shown) we also found out that the overlap between the three input patterns (limited to 10 % by default; see Section 2.2.1) has a large impact on the emergent assemblies. Higher overlaps in the input patterns generally lead to larger numbers of MRUs and smaller assemblies after the encoding phase (and subsequently, to lower numbers of PCUs after the association phase; see Section 3.3).

### 3.3 Emergence of associations

Associations between memory items emerged in the experiments of (Ison et al. 2015) by repeatedly presenting to a human subject combinations of two images from a fixed set of images: a particular face image was shown in front of a particular landscape image. The firing activity of a total of 613 units in the human MTL was recorded before, during, and after this creation of an association. Only those images were used for this pairing protocol for which its presentation had previously caused increased firing of at least one neuron in the MTL from which one recorded. These neurons can be viewed as members of the memory traces that encode the corresponding image or concept. A key finding of Ison et al. (2015) was that the two assemblies that encoded the two components of a combined image changed during the formation of an association between them: each of them expanded, and recruited neurons from the other assembly.

In order to mimic this experimental setup in our model, we repeatedly presented combinations of two of the previously used external input patterns, the blue and the green pattern. This combined pattern was constructed by superimposing (adding) the stationary rate patterns of the 200 external input neurons for the two input patterns, truncated at 40 Hz. The resulting combined input pattern with superimposed noise was presented 20 times at random time points within 36 s of a subsequent “association phase” (see Fig. 2). Figure 4B shows that synaptic weights between neurons in the two associated assemblies were rapidly increased during this phase.

Figures 5A and B illustrate the network activity in response to (A) the first presentation of the combined pattern during the association phase and (B) a presentation of the combined and the three separate input patterns directly after the association phase. We found that in this simulation 9 neurons in the blue assembly 1 (indicated by the green horizontal lines in Figure 5B) became also members of the green assembly. Simultaneously 15 members of the green assembly 2 (indicated by the blue horizontal lines in Figure 5B) became also members of the blue assembly. In analogy to the terminology of (Ison et al. 2015) we refer to these neurons in the resulting overlap of the two assemblies as pair-coding units (PCUs; exact definition in Section 2.3.2). We also say that the blue input pattern is the preferred (“P”) stimulus for neurons in the blue assembly, whereas the green pattern is the non-preferred (“NP”) stimulus for neurons in this assembly (analogous for the green assembly). 21 out of 51 neurons in the data of (Ison et al. 2015) initially preferred one of the two image components (P stimulus), but responded after the association with an increased firing rate also to the non-preferred (NP) stimulus. A very similar scenario emerged in our network model, where 24 out of 54 neurons in the blue and green assembly responded after the association phase with an increased firing rate to the NP stimulus. We will analyze in Section 3.5 how this fraction depends on parameters of the model.

Since the neurons in our model produce spike trains, we can directly compare changes in firing responses of neurons before and after the induction of the association between the data (Figure 5A-C of Ison et al. 2015) and our model (see Figure 5C-E). We find that the neural coding properties of PCUs in the blue and green assemblies change through STDP after repeated presentations of combined input patterns in a way that is very similar to the data of (Ison et al. 2015): The firing response remains significant for the preferred stimulus, changes from insignificant to significant for the non-preferred stimulus, and remains insignificant for the red (“non-associated” or “NA”) input pattern that was not part of the combined stimulus (see Figure 5C-E).

A neuron can become through synaptic plasticity in two different ways a member of the assembly for the previously non-preferred stimulus: By increasing its weights from input neurons that are highly active during the NP stimulus, or by increasing the weights from neurons in the assembly for the NP stimulus. A latency analysis in (Ison et al. 2015) arrived at the conclusion that a combination of both effects occurred. In our model we can measure directly how much synaptic input a neuron that starts to respond to the NP stimulus after the association induction gets from the external input neurons, and how much from the original assembly for the NP stimulus. For that purpose we carried out experiments where (i) all internal connections from the assembly of the NP or (ii) all connections from the input were disabled. The resulting PCU activities in response to the NP stimulus can be found in Figure S5 in the *Supplementary Material*. These results show that also in the model a combination of both effect occured, with the contribution from the external input neurons being somewhat stronger.

### 3.4 Neuronal and functional learning curves

A key point of the experimental data of (Ison et al. 2015) was that the overlap of assemblies emerged at about the same presentation of the combined stimulus when the association between the two memory traces became functional, i.e., when the subject was able to select in an interjected multiple choice test the correct background that had previously been shown in conjunction with a face.

We asked whether the same effect would occur in our model, i.e., whether the overlap between the associated blue and green assembly would emerge at about the same presentation (“trial”) when a downstream network would be able to detect a functional association between the two assemblies, i.e., detect an activation of an assembly when its associated assembly was activated via an external input. In addition, we were able to investigate in the model a question that could not be probed through recordings from the human MTL: will weights of synapses that interconnect neurons in the two assemblies increase significantly through the association process, and will a significant weight increase appear at about the same time as the overlap between the two assemblies? If this is the case, it suggests that, in addition to the emergent overlap, this weight increase is also related to the emergent functionality of the association.

The mechanisms by which downstream networks in the brain extract information from assembly activations in the MTL for decision making are largely unknown. Hence we employed two arguably minimal models for that:

#### Readout A

Integration of evidence by counting spikes in the associated and the non-associated assembly, with a subsequent symbolic multiple choice test, where the assembly with the higher spike count is selected as associated assembly. This multiple choice test can be seen as a simple realization of the multiple choice test in (Ison et al. 2015).

#### Readout B

Linear readout neurons are trained for each of the assemblies to fire whenever this assembly is activated through the corresponding external input. Then test whether the readout neuron for the green assembly signals that the green assembly is activated when the external input for the blue assembly is injected, and vice versa.

The results of (Yang and Shadlen 2007) suggest that competing neurons in the parietal lobe are able to integrate evidence for and against a decision during a time span of over 2 s in their firing rates. Phenomenological their response is not so different from the assumed readout neurons in model A. Model B is based on the assumption that assembly activations themselves are fundamental signals in inter-area communication, so that the activation of a particular assembly in the MTL triggers the firing of a corresponding neuron in downstream networks. Although both readout models are not fully supported by experimental data, they appear to be reasonable choices as simple minimal models.

The results are shown in Figure 6. Solid curves show averages over 20 experiments with random network realizations and input pattern triples. Small circles show results for the standard model. One finds that the functional performance of readout B tracks the evolution of both the overlap between the two assemblies (PCU fraction in A) and mean synaptic weight between neurons in these two assemblies (panel B) very well. Readout A in panel C, which models a multiple choice test, shows good performance already after just one presentation of the combined stimulus.

One should point out that the interleaved tests of associations for the human subjects (interleaved Tasks 2 and 3 in Ison et al. 2015) may have enhanced learning processes beyond the learning from passive presentation as in our model.

### 3.5 Emergence of a hierarchical neural code for associated memory traces

The previously discussed results of the model were based on our standard values for parameters that affect the size of the network response to single and combined external stimuli, and thereby also the impact of STDP for synaptic connections between excitatory neurons. In this way they also affect the number of PRUs, i.e., the number of neurons that become members of an assembly, and the number of PCUs, i.e., the resulting size of the overlap of the two assemblies for which combinations of the corresponding patterns had been presented.

We show here that if one changes two of these parameters, a different neural code emerges for associated memories, namely the hierarchical model in the terminology of (Kahana et al. 2008, Kahana 2012). According to this model neurons emerge, which we call HUs, that are activated by a combined input pattern, but are not activated by either of its components. The relative sizes of the number of HUs and PCUs can be seen as a measure to what extent a hierarchical model or a chaining model is expressed.

Critical parameters for switching between the two competing memory models are *γ*_exc_, which scales the excitability of excitatory neurons, and *γ*_w_, which scales initial weights to excitatory neurons. Figure 7 shows their impact on the mean assembly size and the numbers of MRUs, PCUs, and HUs in 2D surface plots. The black star marks their values in the previously discussed standard model. Intersections of grid lines indicate computed values (9 *×* 9 grid), whereas intermediate grid values were interpolated.

Figure 7D shows a peak value of HUs at *γ*_w_ = 1.0 and *γ*_exc_ = 0.25 (with a low value of PCUs) and a peak value of PCUs at *γ*_w_ = 0.71 and *γ*_exc_ = 0.25 (with a low value of HUs). Hence our model can switch between a hierarchical and a chaining model for the formation of associations by changing these two parameters. The absolute numbers of PCUs or HUs were found to be increasing with increasing values of *γ*_exc_. Figure S6 in the *Supplementary Material* illustrates the network activity in response to different input patterns for the setup with *γ*_w_ = 1.0 and *γ*_exc_ = 0.25, where the hierarchical memory model emerges through STDP (compare with Figure 5, representing the chaining model). No overlap (i.e., no PCUs) between the blue and green assemblies was found in this case.

In order to test whether the network operated for both of these parameter settings in a biologically reasonable regime, we computed the I-to-E ratio 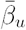 based on membrane potentials at three time points of interest: (i) after the initialization phase, (ii) directly before, and (iii) directly after the association phase. The impact of *γ*_exc_ and *γ*_w_ on the E/I balance is shown in Figure S8 in the *Supplementary Material* for these three time points. A small increase of 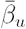 during the course of the simulation can be seen. We found that inhibition dominates excitation for both of the previously discussed ranges of *γ*_exc_ and *γ*_w_, that yield different neural codes for associations. This finding in the model is in agreement with experimental findings in neocortex (Haider et al. 2013) and area CA3 of the hippocampus (Calfa et al. 2015, Atallah and Scanziani 2009). Hence both the parameter values where STDP induces many PCUs, i.e., the chaining model for associations, and where STDP induces many HUs, i.e., the hierarchical model, yield a biologically plausible regime where inhibition dominates excitation.

The impact of other network parameters, such as *E*_exc,generic_, *I*_exc,generic_, and *p*_E*→*E_ is shown in Figure S7.

### 3.6 Mathematical analysis of the formation of an association on assembly representations

The results presented in this article arise from the simulation of a generic neural network model that is as compatible as possible with what we know about the cortex: A recurrent network of stochastically spiking neurons with inhibition, STDP, and short-term plasticity. In this section we address the question: Can we achieve a level of mathematical understanding of the assembly association phenomenon that complements our simulation results?

To our knowledge, there are no known techniques for analyzing mathematically recurrent networks of spiking neurons, especially under biologically realistic STDP and shortterm plasticity, beyond severely simplified dynamical systems for the firing rates such as (Abbott 1994). However, a deeper mathematical understanding of the phenomenon of assembly association is possible, albeit at the expense of neurorealism. In a companion work to the present paper we have advanced a mathematical analysis of the formation and association of neuronal assemblies (Legenstein et al. 2018) in a stylized model in which excitatory neurons fire in synchrony and in discrete time steps, instead of spiking stochastically; Hebbian plasticity replaces STDP, and there is no short-term plasticity; the recurrent and afferent synaptic networks between excitatory neurons constitute an Erdős–Rényi random graph (Erdős and Rényi 1959); finally, explicit inhibition is replaced by a “k winners take all” rule for the firing of excitatory neurons. One can show analytically (Legenstein et al. 2018) that in this simplified model an assembly is formed in response to a stimulus, and that the overlap of two assemblies is increased to reflect association due to the simultaneous presentation of two stimuli, all with high probability (where the synaptic network is the source of randomness). Hence a theoretical analysis for a simplified neural network model corraborates the findings of this paper. Further mathematical insights into assemblies and their interaction –including the formation of hierarchical associations as pointed out in the present paper– are the subject of on-going work.

One further theoretical question raised by the results of (Ison et al. 2015, De Falco et al. 2016) and the present study is whether a realistic web of overlaps between assemblies can be created in the brain to encode complex associations between the corresponding real-life concepts. A natural question in this regard is the following: what are the limitations on the structure of such an “association graph” –whose vertices represent concepts and whose edges represent associations– so that the graph can be realized in cortex through the representation of vertices by assemblies and edges by intersecting assemblies, assumed to be all of the same size? New mathematical results (Anari et al. 2018; Section 4) establish that, rather surprisingly, quite general webs of associations can in principle be encoded through overlaps of relative size similar to those observed experimentally in (Ison et al. 2015, De Falco et al. 2016). The only limitation that arises from this general theoretical analysis concerns the maximum degree of the web of associations, i.e., the maximum number of concepts that can be associated with a given one. In principle a maximum degree of 40 is supported by this analysis. However a biologically more realistic emulation, where assemblies and their intersections emerge from random choices of neurons, arrives at a smaller upper bound of 10 for the degrees of vertices. These theoretical results suggest that implementations of webs of associations in the brain through assemblies and their intersections are necessarily imperfect as far as concepts with substantially more associations are concerned: missing associations to single concepts (possibly alleviated by more reliable associations when further concepts are added to recall the context) or spurious associations are likely to occur.

## 4. Discussion

The emergence of memory traces and associations between memory traces are fundamental for most higher cognitive functions of the brain. Nevertheless, it remains poorly understood how these processes are implemented in neural networks of the brain. Theoretical and modelling analyses are likely to shed light on this problem. We propose that a recurrent network of spiking neurons with data-constrained short-term and long-term synaptic plasticity provides a suitable framework for that. We found that the emergence of memory traces for repeatedly occurring external inputs can be reproduced in such a model through STDP for synaptic connections between excitatory neurons. Furthermore we found that two different coding mechanisms for the formation of associations between memory traces can be reproduced in the model. One coding mechanism relies on the emergence of overlaps between memory traces, as proposed by chaining models (Kahana et al. 2008, Kahana 2012), and supported by recordings from the human MTL (Ison et al. 2015, De Falco et al. 2016). A different neural coding mechanism was proposed by hierarchical models (Kahana et al. 2008, Kahana 2012, Valiant 2000a;b, Norman and O’Reilly 2003), and was supported by recordings from the rodent brain (Komorowski et al. 2009). It postulates that the memory traces themselves remain largely unchanged during the formation of an association between them, and that instead new neurons are recruited for encoding the combined memory.

In a family of more abstract memory models each memory item is represented by a vector of numbers. The combination of two memory items is then postulated to be encoded by an addition, concatenation, or convolution of the two vectors of numbers (Fodor and Pylyshyn 1988, Smolensky 1990, Kanerva 1994, Rizzuto and Kahana 2001, Plate 2003). In a common method for mapping such vectors of numbers to activity patterns in neural networks, see e.g. (Eliasmith 2013), the neural activity that represents the new vector contains many neurons that are not active in the neural representation of either of the two original vectors. This convention makes these abstract models similar to hierarchical models with regard to the question whether new neurons are recruited for the combined memory. Also a model from theoretical computer science (Valiant 2000a;b; 2005) predicts that the neural code for two associated memory items consists of neurons that do not belong to either of the assemblies for the two memory components.

We identified two parameters of our model, which control the excitability of excitatory neurons and the scale of initial synaptic weights between them, as being critical for the resulting type of neural code for associated memories (Fig. 7). Both of these parameters appear to be “soft” parameters, whose values can be changed in an adaptive manner by the brain. The excitability of neurons is known to be regulated on several time scales, see e.g. (Debanne et al. 2019) for a recent review. The initial synaptic weights before a new association is formed is obviously subject to a host of preceding plasticity processes. Hence our model suggests that the brain is able to use both of these two neural codes, and possibly switch between them during consolidation.

On the side our model shows that –in contrast to an assumption of a recently published theoretical model (Legenstein et al. 2018)– the number of connections within an emergent assembly is in our model usually not above average. While the overall recurrent connection probability is fixed, the instantiation of a concrete wiring could be such that the actual number of connections and therefore, the connection probability within an assembly may or may not be higher than average. Indeed, an analysis of recurrent connection probabilities within assemblies for 20 simulations with random realizations and input patterns showed that connection probabilities within these assemblies were higher than within random subsets in just 25 % of the cases (*α* = 5 % Monte-Carlo permutation test with *N* = 1000).

In contrast to the large scale model for the formation of memory traces –without associations between them– in (Guzman et al. 2016), we have investigated here a minimal model with more physiological details for the formation of associations. The next step will be to combine both modelling approaches in a large scale model with neurophysiological details. Such a model is likely to provide theoretical insight into the way how the intricate web of associations is formed and continuously updated in the human brain. Thereby it will help us to understand one of the most amazing information processing capabilities of the human brain.

## Acknowledgements

Written under partial support by the Human Brain Project of the European Union #604102 and #720270, the Austrian Science Fund (FWF): I 3251-N33, and the National Science Foundation of the USA #1408635

This article has been accepted for publication in *Cerebral Cortex* published by Oxford University Press.

## Supplementary Material

### Generation of memory traces

**Figure S1:**
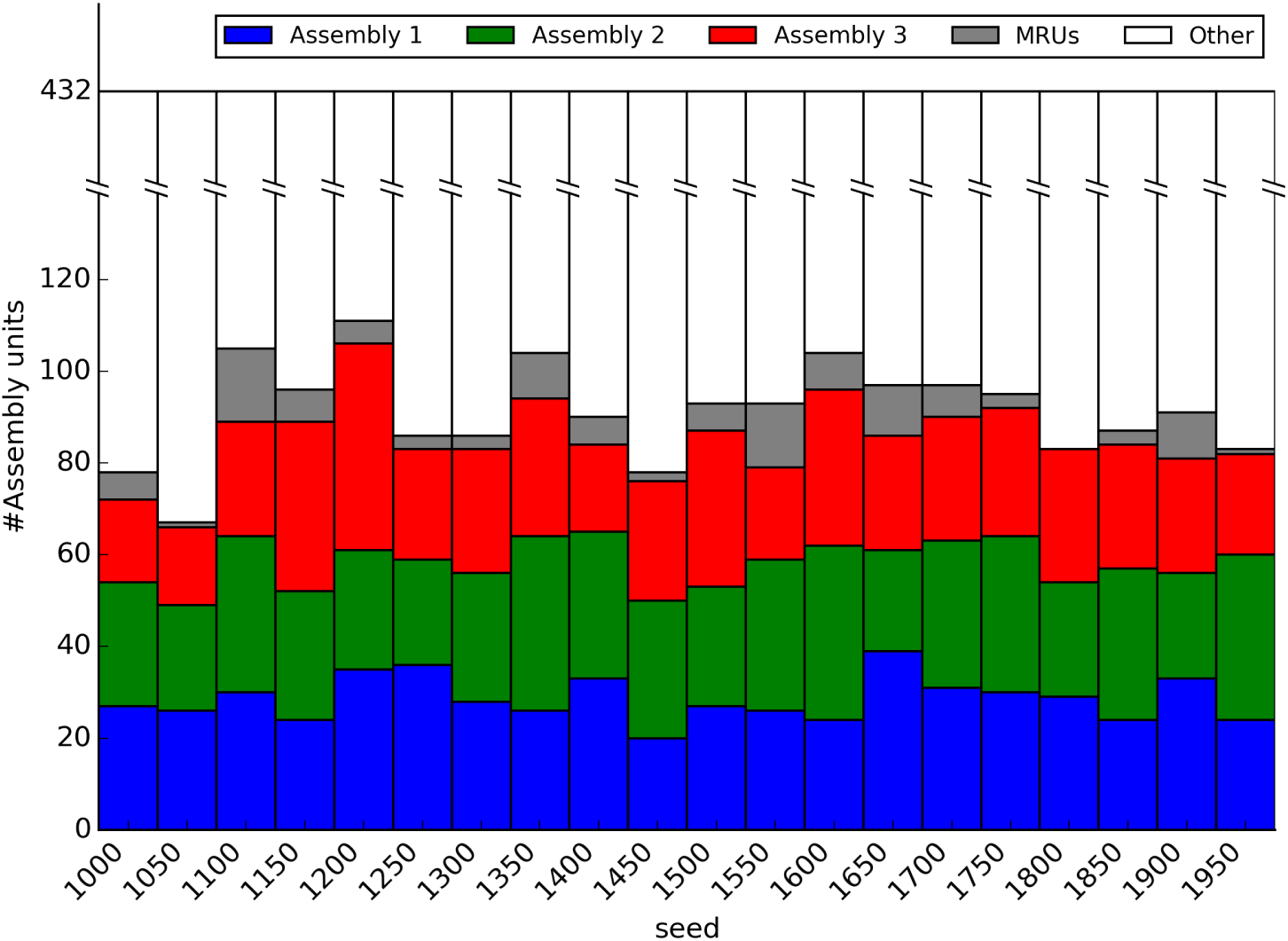
Overview of assembly sizes (blue, green, red) and MRUs (yellow) over 20 simulations with different random seeds.

**Figure S2:**
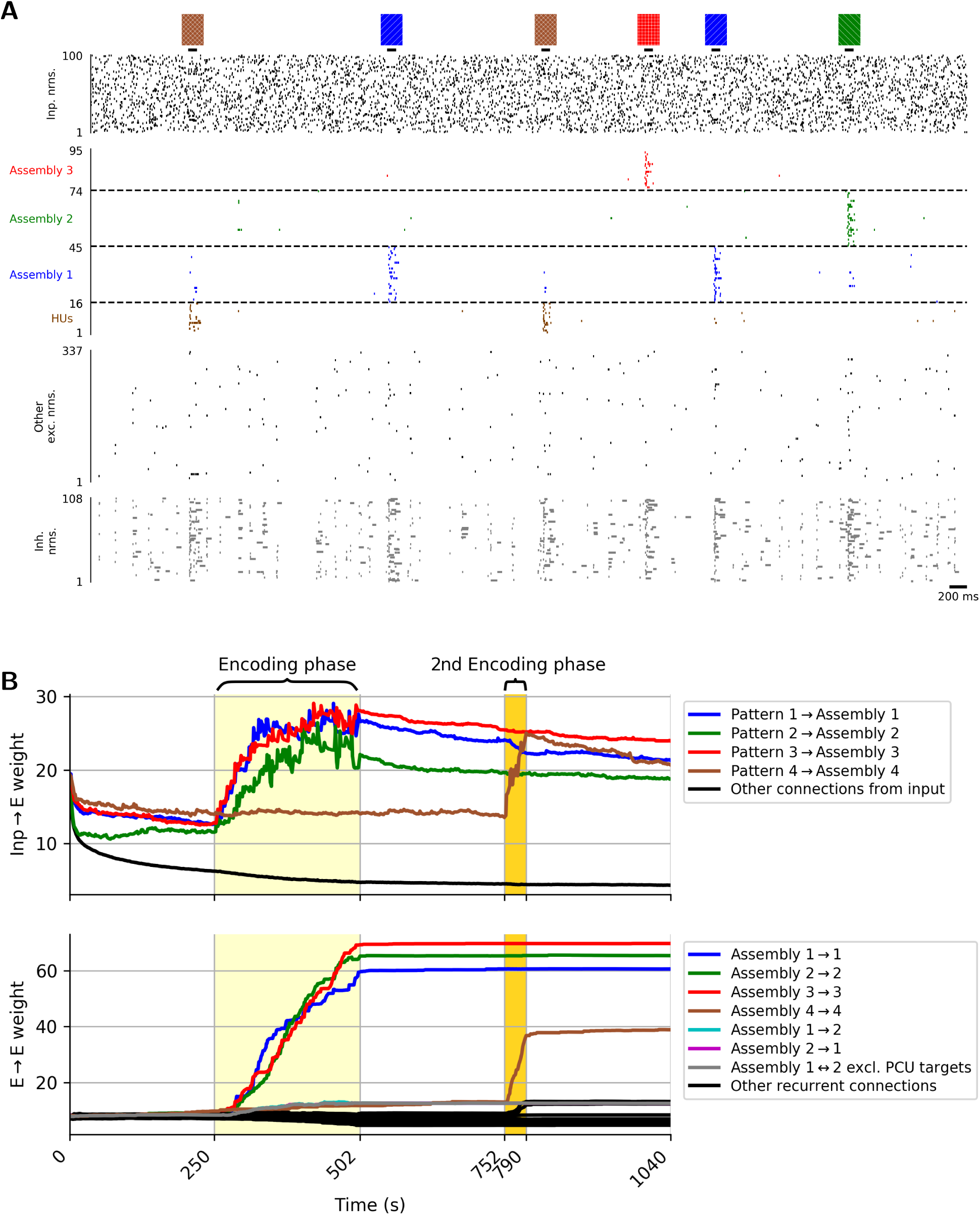
(A) Emergence of a fourth assembly (brown color) after a new input pattern has been introduced during a 2nd encoding phase (instead of combinations of previously introduced patterns). (B) Mean weight changes of connections from input neurons to the resulting four assemblies (taking only on-channels of the stationary input rate patterns into account) and internal connections within the four assemblies during all phases of the main simulation.

### Emergence of associations

**Figure S3:**
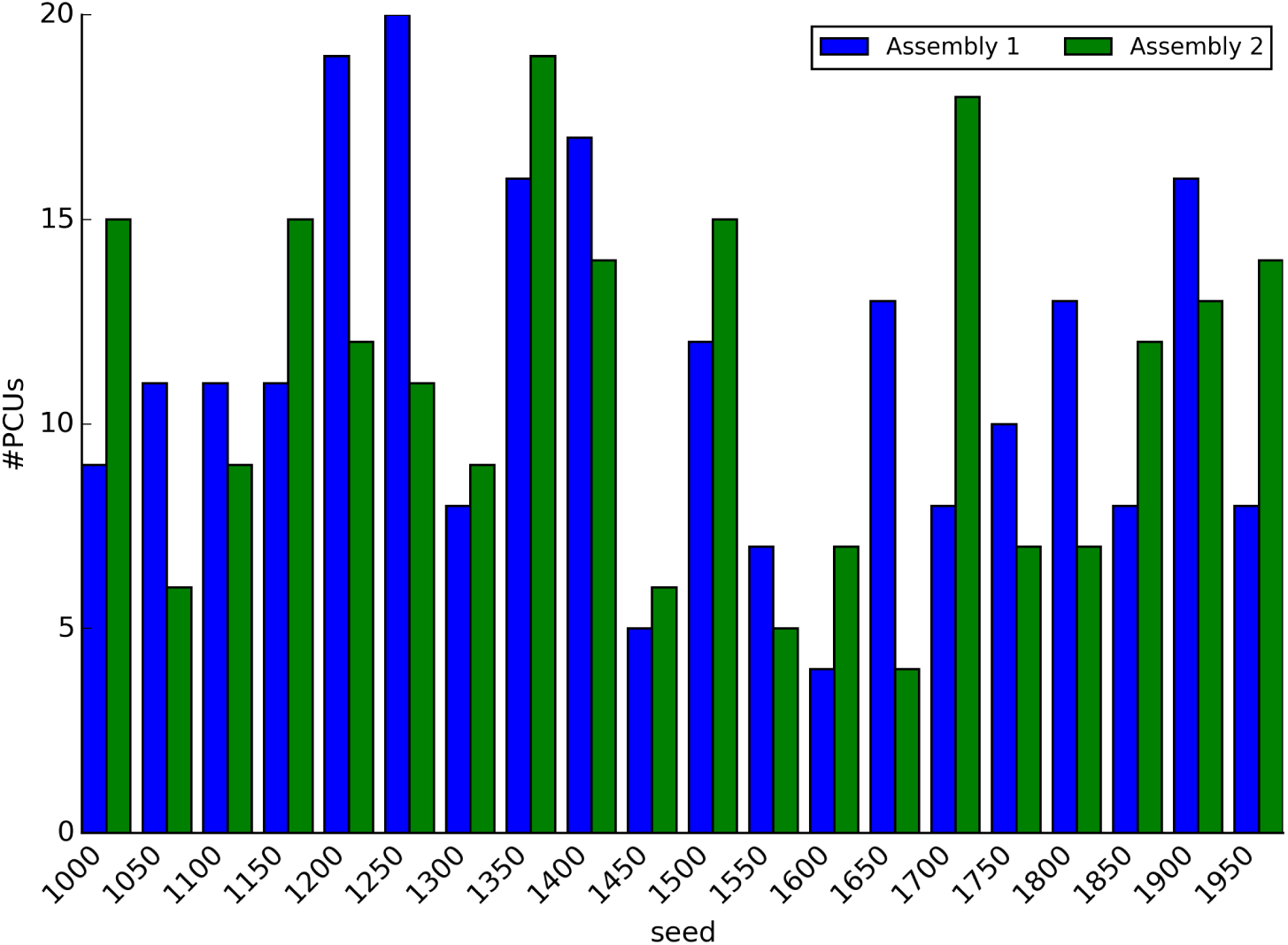
Overview of the numbers of PCUs per (blue and green) assembly over 20 simulations with different random seeds.

**Figure S4:**
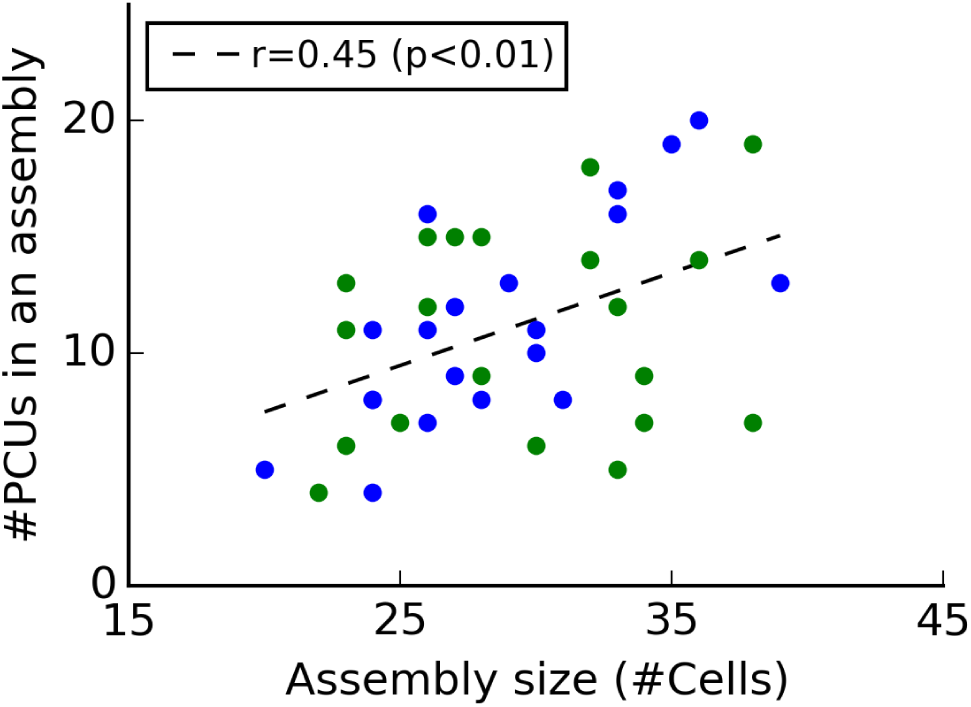
Scatter plot showing a weak but significant linear correlation between the assembly sizes and the resulting numbers of PCUs per (blue and green) assembly over 20 simulations (*r* = 0.45, *p <* 0.01).

**Figure S5:**
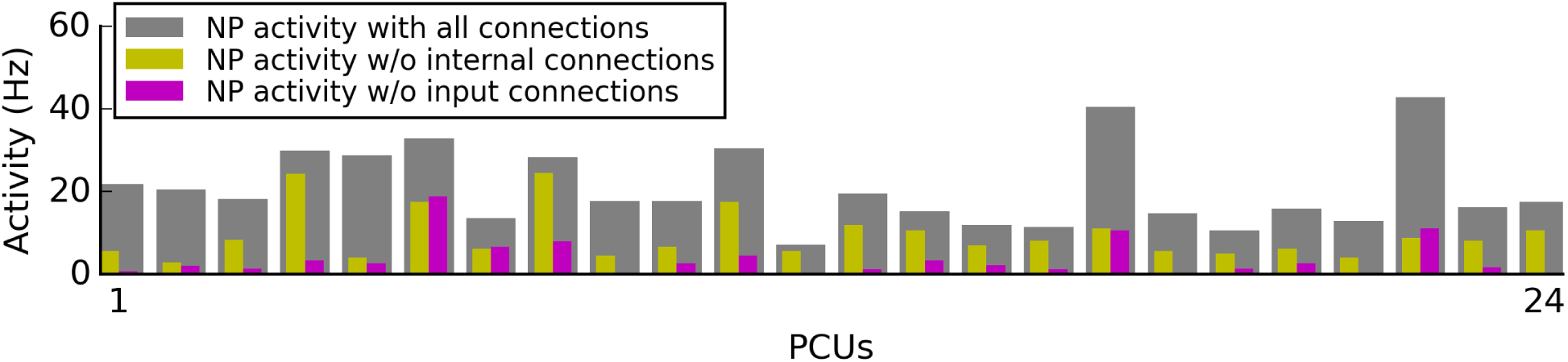
Sources of synaptic inputs to PCUs for the NP stimulus. Grey bars: Firing rates of PCUs with all connections intact. Yellow bars: Firing rates after the internal connections from the assembly of the NP stimulus were disabled. Magenta bars: Firing rates after the input connections were disabled.

### Functional impact of network parameters

**Figure S6:**
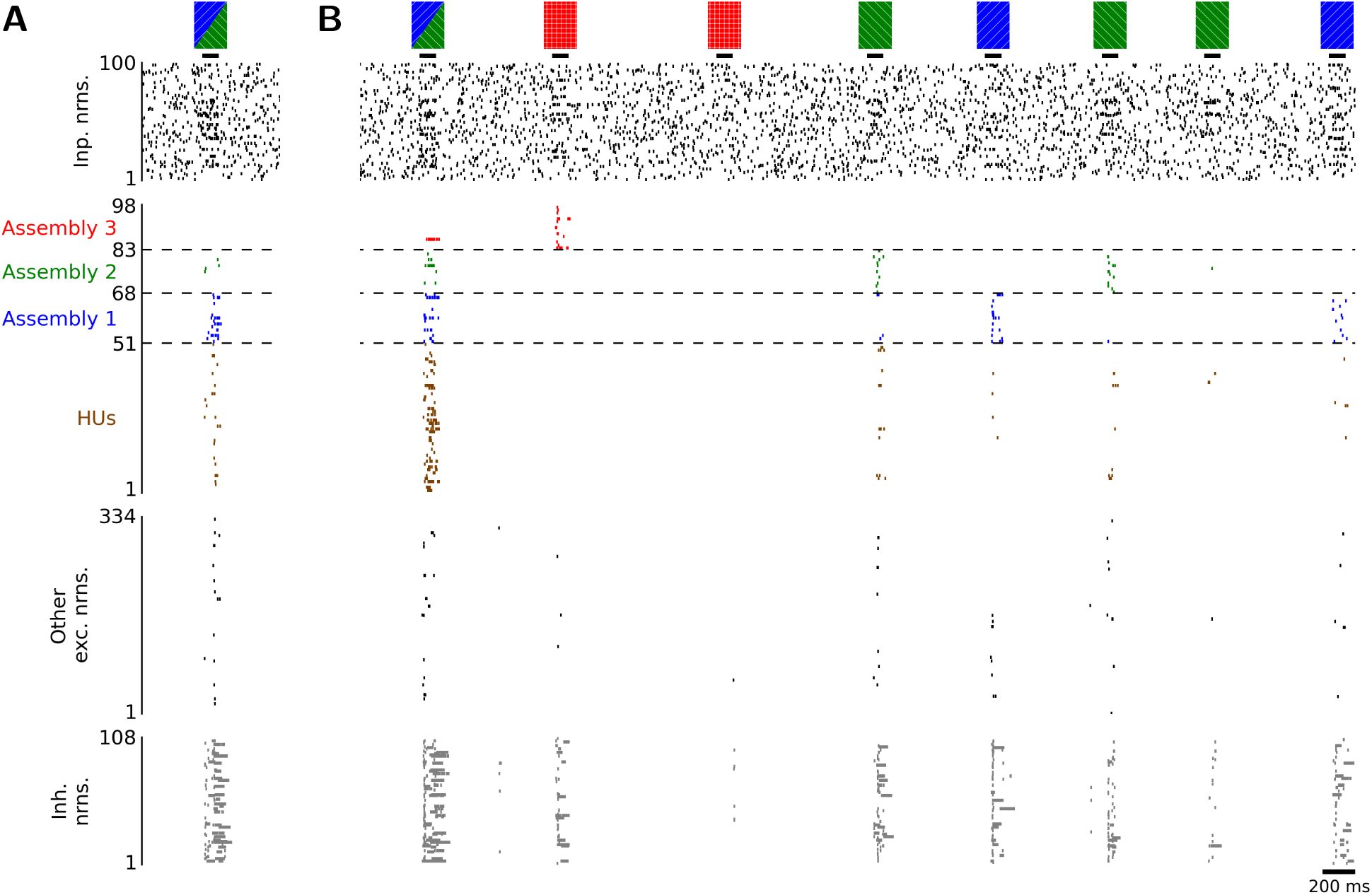
HUs as emergent neural code for associations, shown for a network with *γ*_w_ = 1.0 and *γ*_exc_ = 0.25 (compare with Figure 5). (A) Response of the network to the first presentation of such a combined pattern during the association phase. (B) After 20 presentations of this combined input pattern during the association phase, a new memory trace represented by 51 HUs encoding the combined input patterns, instead of an overlap between the blue and green assemblies (i.e., no PCUs), emerged.

**Figure S7:**
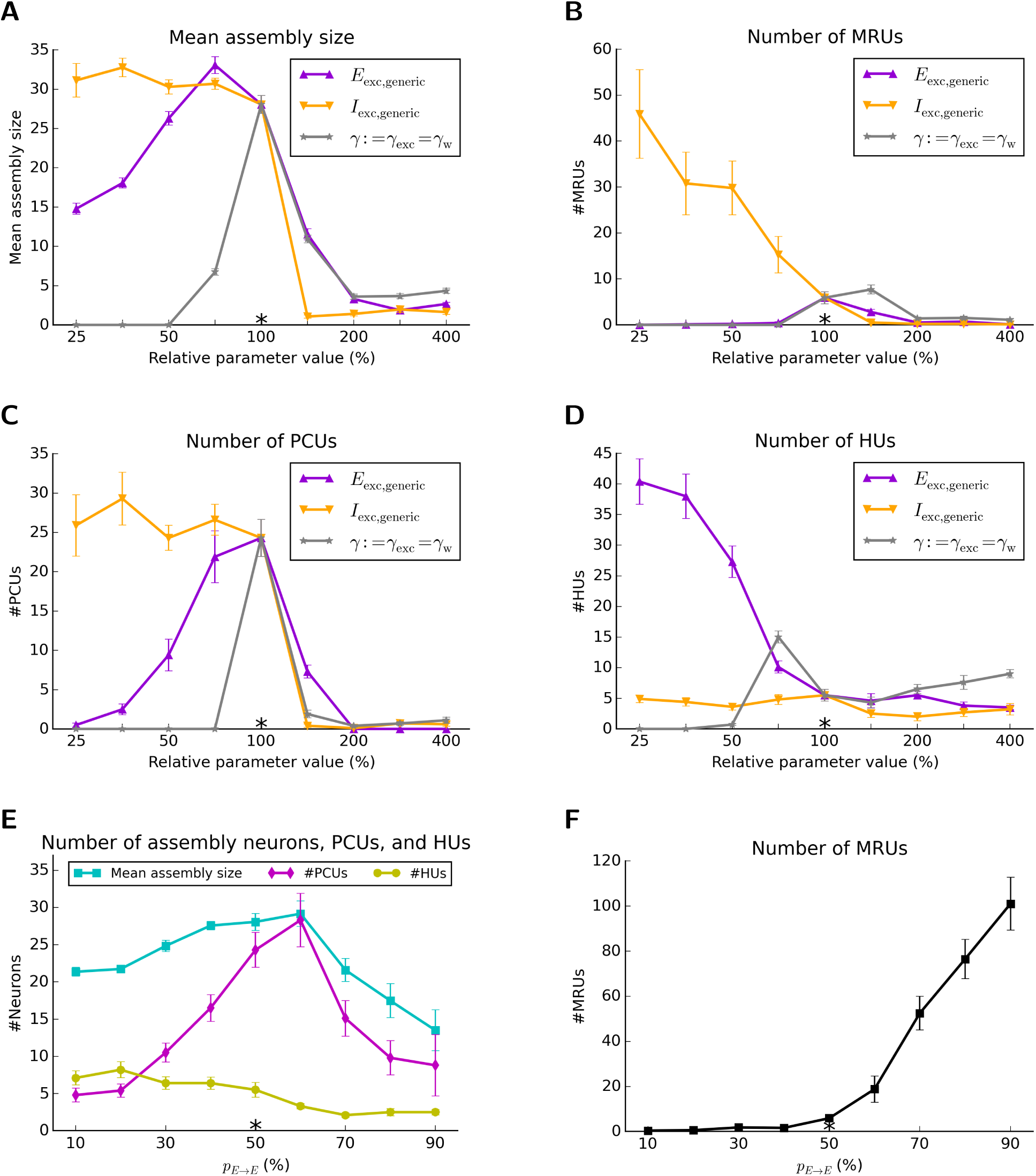
Impact of various network parameters on the mean assembly size, and the numbers of MRUs, PCUs, and HUs. Mean values were estimated over 10 simulations with different random seeds. Error bars represent the SEM. The * symbols mark the standard parameter values. (A-D) The parameter values of *E*_exc,generic_, *I*_exc,generic_, and *γ* := *γ*_exc_ = *γ*_w_ were varied in logarithmic steps from 25 % to 400 % relative to the respective standard values. (E-F) The recurrent excitatory connection probability *p*_E*→*E_ was varied in linear steps from 10 % to 90 %. More details can be found in the supplemental text below.

In Figure S7A-D, the dependence of the mean assembly size and the numbers of MRUs, PCUs, and HUs on three parameters was investigated in logarithmic steps: *E*_exc,generic_ regulates the generic excitability of the excitatory population, *I*_exc,generic_ regulates the generic excitability of the inhinitory population, and *γ* scales the relative contributions of the generic excitability of the excitatory population and the initial synaptic weights between Inp*→*E and E*→*E connections. We found that the mean assembly size and the number of PCUs roughly peak for values close to the standard value of *γ* and *E*_exc,generic_ while they are constantly low for *I*_exc,generic_ larger than the standard value and constantly large otherwise. The number of MRUs is negatively correlated with *I*_exc,generic_ while being constantly low and largely unaffected by the two other parameters. The number of HUs was found to be negatively correlated with *E*_exc,generic_ while being constantly low and largely unaffected by the two other parameters, with a small peak at *γ* = 70.7 %.

In Figure S7E-F, we investigated the impact of the connection probability *p*_E*→*E_ between pairs of excitatory neurons in the recurrent network, using again standard values for all other parameters. The mean assembly size and number of PCUs were found to peak at a connection probability of *p*_E*→*E_ = 60 % whereas the number of HUs is largely unaffected by the connection probability. The number of MRUs is highly increasing with increasing levels of *p*_E*→*E_. A proper choice of this connection probability is difficult for a model, because its impact on the number of PCUs depends on the size of the neural network model. From the functional perspective it is essential from how many neurons in the same and the associated assembly a neuron in an assembly receives synaptic connections. If this number, which depends on the connection probability and the network size, is too low, few PCUs are likely to emerge. Replacing the biologically more realistic synapse models with short-term plasticity by simpler synapse models with static efficacies did not affect the main results of this study.

A simple calculation shows the following for the subarea CA3 of the hippocampus, which is estimated to consist of 2.83 million pyramidal cells in humans (Andersen et al. 2007) and has an estimated connection probability between pyramidal cells of 0.92 % in rodents (Guzman et al. 2016): An assembly for a memory item in the human MTL was estimated to consist of between 0.2 and 1 % of the pyramidal cells in the MTL (Waydo et al. 2006). If one assumes that this also holds for area CA3, one arrives at an estimate of 5660 to 28300 for the number of neurons in an assembly in area CA3. Thus with a connection probability of 0.92 %, each neuron in one assembly receives on average synaptic input from 52 to 260 neurons in any other assembly. This suggests that models of different network sizes should have a connection probability that scales this number of presynaptic neurons from another assembly into a comparable range, so that they can contribute significantly to its firing probability. In our small neural network model this number of presynaptic neurons from another assembly had an average value of 12, but in order to achieve that we had to increase the connection probability between excitatory neurons to an unrealistically large value of 50 %.

### Balance between excitation and inhibition

**Figure S8:**
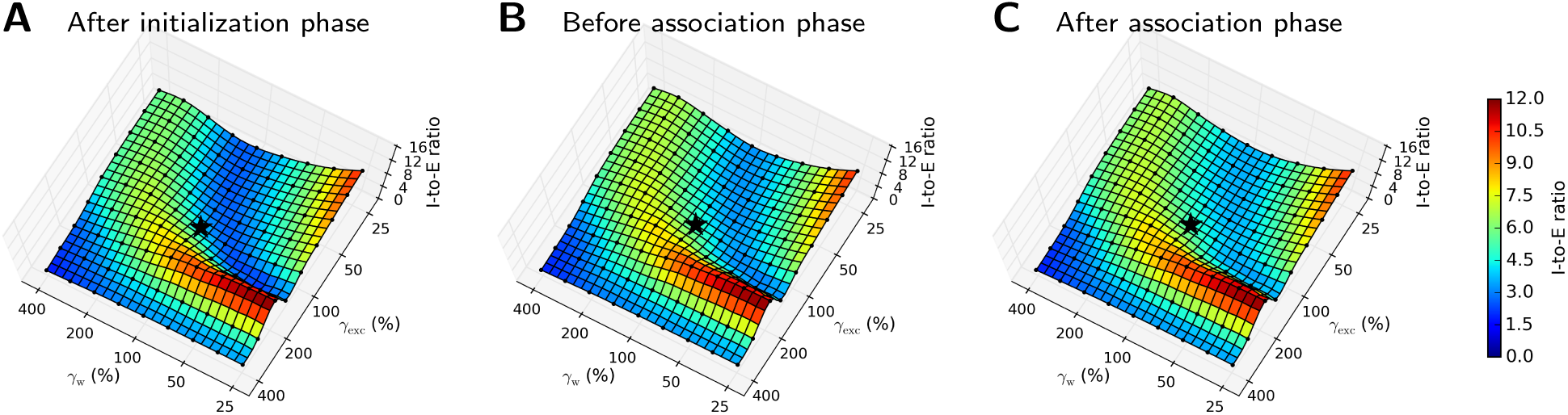
Impact of the scaling factors *γ*_exc_ and *γ*_w_ on the E/I balance based on membrane potentials at time points (A) after the initialization phase, and directly (B) before and (C) after the formation of associations. Mean values were estimated over 10 simulations with different random seeds. The * symbols mark the standard parameter values. The parameter values were independently varied in logarithmic steps from 25 % to 400 % relative to the respective standard values, as indicated by the black dots. Intermediate grid values were interpolated.

## References

L. Abbott. Decoding neuronal firing and modelling neural networks. Quarterly Reviews of Biophysics, 27(3):291–331, 1994.

N. Anari, C. Daskalakis, W. Maass, C. Papadimitriou, A. Saberi, and S. Vempala. Smoothed analysis of discrete tensor decomposition and assemblies of neurons. In S. Bengio, H. Wallach, H. Larochelle, K. Grauman, N. Cesa-Bianchi, and R. Garnett, editors, Advances in Neural Information Processing Systems 31, pages 10880–10890. 2018.

P. Andersen, R. Morris, D. Amaral, T. Bliss, and J. O’Keefe. The Hippocampus Book. Oxford University Press, 2007.

B. V. Atallah and M. Scanziani. Instantaneous modulation of gamma oscillation frequency by balancing excitation with inhibition. Neuron, 62(4):566–577, 2009.

G. Buzsáki. Neural syntax: cell assemblies, synapsembles, and readers. Neuron, 68(3): 362–385, 2010.

G. Calfa, W. Li, J. M. Rutherford, and L. Pozzo-Miller. Excitation/inhibition imbalance and impaired synaptic inhibition in hippocampal area CA3 of Mecp2 knockout mice. Hippocampus, 25(2):159–168, 2015.

E. De Falco, M. J. Ison, I. Fried, and R. Quian Quiroga. Long-term coding of personal and universal associations underlying the memory web in the human brain. Nature Communications, 7:13408, 2016.

D. Debanne, Y. Inglebert, and M. Russier. Plasticity of intrinsic neuronal excitability. Current Opinion in Neurobiology, 54:73–82, 2019.

C. Eliasmith. How to build a brain: A neural architecture for biological cognition. Oxford University Press, 2013.

J. Eppler, M. Helias, E. Muller, M. Diesmann, and M.-O. Gewaltig. PyNEST: A convenient interface to the NEST simulator. Frontiers in Neuroinformatics, 2(12), 2009.

P. Erdős and A. Rényi. On random graphs. Publicationes Mathematicate, 6:290–297, 1959.

J. A. Fodor and Z. W. Pylyshyn. Connectionism and cognitive architecture: A critical analysis. Cognition, 28(1):3–71, 1988.

A. Gupta, Y. Wang, and H. Markram. Organizing principles for a diversity of GABAergic interneurons and synapses in the neocortex. Science, 287(5451):273–278, 2000.

S. J. Guzman, A. Schlögl, M. Frotscher, and P. Jonas. Synaptic mechanisms of pattern completion in the hippocampal CA3 network. Science, 353(6304):1117–1123, 2016.

B. Haider, M. Häusser, and M. Carandini. Inhibition dominates sensory responses in awake cortex. Nature, 493(7430):97, 2013.

K. L. Hoffman and N. K. Logothetis. Cortical mechanisms of sensory learning and object recognition. Philosophical Transactions of the Royal Society of London B: Biological Sciences, 364(1515):321–329, 2009.

M. J. Ison, R. Quian Quiroga, and I. Fried. Rapid encoding of new memories by individual neurons in the human brain. Neuron, 87(1):220–230, 2015.

R. Jolivet, A. Rauch, H.-R. Lüscher, and W. Gerstner. Predicting spike timing of neocortical pyramidal neurons by simple threshold models. Journal of Computational Neuroscience, 21(1):35–49, 2006.

R.S.A. Josselyn, S. Köhler, and P. W. Frankland. Finding the engram. Nature Reviews Neuroscience, 16(9):521, 2015.

M. J. Kahana. Foundations of Human Memory. Oxford University Press, 2012.

M. J. Kahana, M. W. Howard, and S. M. Polyn. Associative retrieval processes in episodic memory. semanticscholar.org, 2008.

P. Kanerva. The spatter code for encoding concepts at many levels. In ICANN’94, Springer, London (226–229), 1994.

S. Klampfl and W. Maass. Emergence of dynamic memory traces in cortical microcircuit models through STDP. The Journal of Neuroscience, 33(28):11515–11529, 2013.

R.W. Komorowski, J. R. Manns, and H. Eichenbaum. Robust conjunctive item–place coding by hippocampal neurons parallels learning what happens where. Journal of Neuroscience, 29(31):9918–9929, 2009.

S. Kunkel, A. Morrison, P. Weidel, J. M. Eppler, A. Sinha, W. Schenck, M. Schmidt, S. B. Vennemo, J. Jordan, A. Peyser, et al. NEST 2.12.0, Mar. 2017.URL https://doi.org/10.5281/zenodo.259534.

R. Legenstein, W. Maass, C. H. Papadimitriou, and S. S. Vempala. Long term memory and the densest k-subgraph problem. In Proc. of Innovations in Theoretical Computer Science (ITCS), number 57, 2018.

S. Lim, J. L. McKee, L. Woloszyn, Y. Amit, D. J. Freedman, D. L. Sheinberg, and N. Brunel. Inferring learning rules from distributions of firing rates in cortical neurons. Nature Neuroscience, 18(12):1804, 2015.

LinearSVC. API documentation of scikit-learn 0.18: sklearn.svm.LinearSVC, 2016. URL http://scikit-learn.org/0.18/modules/generated/sklearn.svm.LinearSVC.html. Accessed: 2018-03-02.

A. Litwin-Kumar and B. Doiron. Formation and maintenance of neuronal assemblies through synaptic plasticity. Nature Communications, 5(5319), 2014.

H. Markram, Y. Wang, and M. Tsodyks. Differential signaling via the same axon of neocortical pyramidal neurons. Proceedings of the National Academy of Sciences, 95 (9):5323–5328, 1998.

H. Markram, E. Muller, S. Ramaswamy, M. W. Reimann, M. Abdellah, C. A. Sanchez, A. Ailamaki, L. Alonso-Nanclares, N. Antille, S. Arsever, et al. Reconstruction and simulation of neocortical microcircuitry. Cell, 163(2):456–492, 2015.

K. A. Norman and R. C. O’Reilly. Modeling hippocampal and neocortical contributions to recognition memory: a complementary-learning-systems approach. Psychological Review, 110(4):611, 2003.

M. Okun and I. Lampl. Balance of excitation and inhibition. Scholarpedia, 4(8):7467, 2009.

F. Pedregosa, G. Varoquaux, A. Gramfort, V. Michel, B. Thirion, O. Grisel, M. Blondel, P. Prettenhofer, R. Weiss, V. Dubourg, J. Vanderplas, A. Passos, D. Cournapeau, M. Brucher, M. Perrot, and E. Duchesnay. Scikit-learn: Machine learning in Python. Journal of Machine Learning Research, 12:2825–2830, 2011.

J.-P. Pfister and W. Gerstner. Triplets of spikes in a model of spike timing-dependent plasticity. Journal of Neuroscience, 26(38):9673–9682, 2006.

T. A. Plate. Holographic Reduced Representation: Distributed Representation for Cognitive Structures. Stanford, CA, 2003.

R. Quian Quiroga. Neuronal codes for visual perception and memory. Neuropsychologia, 83:227–241, 2016.

D. S. Rizzuto and M. J. Kahana. An autoassociative neural network model of paired-associate learning. Neural Computation, 13:2075–2092, 2001.

P. J. Sjöström, G. G. Turrigiano, and S. B. Nelson. Rate, timing, and cooperativity jointly determine cortical synaptic plasticity. Neuron, 32:1149–1164, 2001.

P. Smolensky. Tensor product variable binding and the representation of symbolic structures in connectionist systems. Artificial Intelligence, 46(1–2):159–216, 1990.

D. Sussillo, T. Toyoizumi, and W. Maass. Self-tuning of neural circuits through shortterm synaptic plasticity. Journal of Neurophysiology, 97(6):4079–4095, 2007.

G. Testa-Silva, M. B. Verhoog, D. Linaro, C. P. De Kock, J. C. Baayen, R. M. Meredith, C. I. De Zeeuw, M. Giugliano, and H. D. Mansvelder. High bandwidth synaptic communication and frequency tracking in human neocortex. PLOS Biology, 12(11): e1002007, 2014.

L. G. Valiant. A neuroidal architecture for cognitive computation. Journal of the ACM, 47(5):854–882, 2000a.

L. G. Valiant. Circuits of the Mind. Oxford University Press, 2000b.

L. G. Valiant. Memorization and association on a realistic neural model. Neural Computation, 17(3):527–555, 2005.

H.-X. Wang, R. C. Gerkin, D. W. Nauen, and G.-Q. Bi. Coactivation and timingdependent integration of synaptic potentiation and depression. Nature Neuroscience, 8(2):187–193, 2005.

S. Waydo, A. Kraskov, R. Quian Quiroga, I. Fried, and C. Koch. Sparse representation in the human medial temporal lobe. The Journal of Neuroscience, 26(40):10232–10234, 2006.

T. Yang and M. N. Shadlen. Probabilistic reasoning by neurons. Nature, 447(7148): 1075, 2007.

F. Zenke, E. J. Agnes, and W. Gerstner. Diverse synaptic plasticity mechanisms orchestrated to form and retrieve memories in spiking neural networks. Nature Communications, 6(6922):1–13, 2015.

